# Explicit Modeling of RNA Stability Improves Large-Scale Inference of Transcription Regulation

**DOI:** 10.1101/104885

**Authors:** Konstantine Tchourine, Christine Vogel, Richard Bonneau

## Abstract

Inference of eukaryotic transcription regulatory networks remains challenging due to the large number of regulators, combinatorial interactions, and redundant pathways. Even in the model system *Saccharomyces cerevisiae*, inference has performed poorly. Most existing inference algorithms ignore crucial regulatory components, like RNA stability and post-transcriptional modulation of regulators. Here we demonstrate that explicitly modeling transcription factor activity and RNA half-lives during inference of a genome-wide transcription regulatory network in yeast not only advances prediction performance, but also produces new insights into gene-and condition-specific variation of RNA stability. We curated a high quality gold standard reference network that we use for priors on network structure and model validation. We incorporate variation of RNA half-lives into the *Inferelator* inference framework, and show improved performance over previously described algorithms and over implementations of the algorithm that do not model RNA degradation. We recapitulate known condition-and gene-specific trends in RNA half-lives, and make new predictions about RNA half-lives that are confirmed by experimental data.

## 1 Introduction

Inference of large-scale transcriptional regulatory networks (TRN) is an active research area, with broad applications in basic science and biomedical research. The key underlying assumption of TRN inference is that changes in RNA expression levels are informative of regulatory relationships between transcription factors (TFs) and their target genes. Information from expression data is often complemented with orthogonal data on protein-protein and protein-DNA interactions, such as from protein binding assays (Valouev et al., 2008; Mundade et al., 2014), DNA accessibility assays like FAIRE-Seq and ATAC-seq (Davie et al., 2015), and motif enrichment analysis (Setty et al., 2015; Guo et al., 2012). Various machine learning approaches are used to infer regulatory networks. They have multiple levels of model complexity, ranging from the earliest Boolean network and network module approaches (Shmulevich et al., 2002; Lähdesmäki et al., 2003; Segal et al., 2003; Pe’er et al., 2001) to approaches that explicitly model dynamics, TF interactions and transcription factor post-transcriptional activity (Honkela et al., 2010; Äijä et al., 2013; Intosalmi et al., 2016; Studham et al., 2014). Advancements in genomics and transcriptomics technologies spurred the development of more complex methods, involving Mutual Information (Margolin et al., 2006a,b; Faith et al., 2007; Butte & Kohane, 2000), correlation (Butte & Kohane, 2000), ANOVA (Küffner et al., 2012), conditional entropy (Karlebach & Shamir, 2012), Random Forest (Huynh-Thu et al., 2010; Petralia et al., 2015), Bayesian causality (Mani & Cooper, 2004; Mani et al., 2012; Friedman et al., 2000), expression module clustering (Reiss et al., 2006, 2015), and constrained regression of biophysical models (Bonneau et al., 2006; Greenfield et al., 2013; Arrieta-Ortiz et al., 2015).

Importantly, recent comprehensive blind assessments of these approaches, called DREAM4 and DREAM5, concluded that there is no single machine learning category of methods outperforming all others (Marbach et al., 2012), and that inference in eukaryotes is systematically much more challenging than in prokaryotes. For example, despite the abundance of data, prediction in yeast *Saccharomyces cerevisiae* only reaches an Area Under Precision-Recall curve (AUPR) of 0:025 and fluctuates around random performance, compared to 0.05 to 0.1 in the bacterium *Escherichia coli*, in which it exceeded random performance several-fold. While DREAM challenges are limited because they did not allow methods to include prior interaction data, results from recent studies that incorporated prior interaction data also dramatically differed between prokaryotes and eukaryotes (Greenfield et al., 2013; Wilkins et al., 2016; Siahpirani & Roy, 2016; Bonneau & Aijo, 2016).

The challenges in inference of eukaryotic networks lie in extensive post-transcriptional regulation. For example, the eukaryotic genome encodes many transcription factors, complex promoter regions, and redundant regulatory pathways. Eukaryotes post-process their RNA and have extensive RNA degradation regulation. However, most inference methods, such as random forest (Huynh-Thu et al., 2010; Petralia et al., 2015), mutual information and related transfer entropy (Margolin et al., 2006a,b), and correlation (Butte & Kohane, 2000), do not explicitly model biophysical parameters involved in expression regulation, and thereby cannot address the increased complexity of eukaryotic expression regulation in a straightforward manner.

The *Inferelator* is a method based on constrained regression (Bonneau et al., 2006; Greenfield et al., 2013). It is distinct from other large-scale inference methods as it allows explicit modeling of biophysical parameters (Equation 1). However, we and others have recently shown that inference of transcription-and translation-related biophysical parameters via ordinary differential equations produces robust genome-wide models of expression changes in response to various stresses in various organisms (Tchourine et al., 2014; Schwanhäusser et al., 2013; Peshkin et al., 2015). Recent modi cations to the Inferelator algorithm, such as robust incorporation of priors (Greenfield et al., 2013) and condition-specific TF activity (TFA) estimation (Liao et al., 2003; Arrieta-Ortiz et al., 2015) dramatically improved the *Inferelator* performance in prokaryotes, boosting the Inferelator performance in *B*. *subtilis* from 0.1 to 0.48 in terms of AUPR. The effect of these modifications in eukaryotic data sets has only been tested in rice (*Oriza sativa*), with modest results (Wilkins et al., 2016)-highlighting the need for new developments that include parameters that account for additional regulatory pathways.

One such crucial regulatory component that is typically ignored (or convolved with other parameters) is RNA degradation. While traditional RNA half-life measurements involved transcription inhibition (Wang et al., 2002; Grigull et al., 2004; Shalem et al., 2008), newer approaches use non-invasive metabolic labeling (Miller et al., 2011; Schwalb et al., 2012; Neymotin et al., 2014). For yeast, several experimental datasets have emerged recently, highlighting the large range in RNA half-lives and their extensive regulation under different conditions and in different genetic backgrounds (Miller et al., 2011; Schwalb et al., 2012; Sun et al., 2012; Neymotin et al., 2014; Munchel et al., 2011). For example, typical RNA half-lives in yeast range between 10 and 18 minutes, while ribosomal genes are twice as long-lived (Neymotin et al., 2014; Munchel et al., 2011; Miller et al., 2011). However, under glucose starvation or rapamycin stress, this relationship becomes inverted, and RNA half-lives systematically stabilize, except for those of ribosomal genes, which become less stable (Munchel et al., 2011). In addition, evidence suggests extensive feedback between transcription and degradation regulation: RNA degradation rates across 46 mutant yeast strains had a strong positive correlation with transcription rates (Sun et al., 2013).

Here, we present the first TRN inference approach that explicitly models RNA half-lives and demonstrate its ability to significantly improve prediction performance. To achieve this result, we first constructed a new, high-quality gold standard of signed transcription regulatory interactions. Then we clustered the 2,577 expression data sets into 20 bi-clusters and showed that combining individually inferred networks outperforms global modeling. We optimized the RNA degradation term for each bi-cluster and showed that the incorporation of this step into modeling further improved performance. Finally, we showed that optimizing network inference for each bi-cluster also results in accurate condition-and gene-specific RNA decay rate predictions. Our final prediction has an AUPR of 0.328, far larger than in other existing work in yeast.

## 2 Materials and Methods

### 2.1 Data Acquisition and Normalization

We acquired four prior known regulatory interaction data sets from various sources, as listed in Table S1, originating predominantly from ChIP-chip, ChIP-seq, knock-out, and overexpression assays (Teixeira et al., 2006; Monteiro et al., 2008; Abdulrehman et al., 2011; Teixeira et al., 2014; Cherry et al., 2012; Costanzo et al., 2014; Kemmeren et al., 2014). The list of 563 TFs was assembled by including all gene names that were marked as either “DNA-binding” or “Regulation of transcription, DNA-templated” in Saccharomyces Genomes Database (SGD) (Cherry et al., 2012; Costanzo et al., 2014), as well as all regulators in the YEASTRACT database of regulatory interactions (Teixeira et al., 2006; Monteiro et al., 2008; Abdulrehman et al., 2011; Teixeira et al., 2014). We downloaded 179 RNA expression data sets from 119 different labs, from Gene Expression Omnibus (GEO) (Edgar et al., 2002; Barrett et al., 2013) using the R Bioconductor package GEOquery (Huber et al., 2015; Davis & Meltzer, 2007). To obtain a high-quality, consistent data set, and to avoid platform-specific batch effects, we exclusively used the Yeast Affymetrix 2.0 platform (GPL2529) for the analysis, as it contained the largest number of unique samples in the GEO database. Raw CEL files for every sample (GSM) measured on this platform were downloaded on March 23, 2015, along with the meta-data for each GSM. We processed and normalized the raw CEL files using the R packages affy (Gautier et al., 2004) and gcrma (Wu et al., 2004), adjusting for background intensities, optical noise, and non-specific binding in a probe sequence specific fashion. The meta-data was processed manually to identify samples that belonged to time series experiments. The full meta-data, as downloaded from the GEO website, is included in the Supplementary File SuppData1.zip. The final RNA expression data set included 2,577 samples, each containing the expression data for 5,716 genes.

### 2.2 Expression Data Clustering

The expression data was further scaled such that every row (gene) had mean 0 and variance 1. The 2,577 expression samples were then clustered using the Euclidean distance metric in R programming language. We first performed PCA on the entire RNA expression matrix, and removed all but the first 16 dimensions to facilitate condition-wise clustering and remove the cumulative effect of noisy low-variance components. We then performed k-means clustering with k = 4. All downstream analysis was performed on the resulting clusters using the original (normalized but unscaled) expression data. The number of clusters was optimized as described in Section 6.1.2.

To annotate the four condition clusters, we used meta-data that we downloaded from GEO for each sample to determine common biological themes, employing the R packages tm (Feinerer & Hornik, 2015; Feinerer et al., 2008) and SnowballC (Bouchet-Valat, 2014). First, we used the binomial test to determine which words are enriched in a given condition cluster as compared to the remaining clusters. To avoid words with a p-value of 0 and minimize lab-specific effects, we then excluded words that had zero counts in all but one cluster. This resulted in a list of words sorted by p-value enrichment in each cluster. p-values were then corrected for multiple hypothesis testing using the Bonferroni correction. Word clouds were created from terms with p-values smaller than 10^−20^, using the wordcloud package in R (Fellows, 2012). The final label assignments were confirmed by manual inspection. Section 6.1.2 provides more details. The list of terms with corrected p-values for each condition cluster and the results of the manual inspection can be found in Supplementary File SuppData2.zip.

In addition to condition-wise clustering, we also performed row (gene-wise) clustering. We first hierarchically clustered the 997 genes in the Gold Standard, and then generalized these clusters to the 5,716 genes present in the entire expression dataset. This resulted in five clusters, for which we performed gene ontology enrichment analysis, as described in Section 2.7. See Section 6.1.3 for more detail.

### 2.3 Curation of the Gold Standard of Regulatory Interactions

A key aspect of the work was the construction of a high-quality gold standard of regulatory interactions in yeast, which was used as prior interactions data for TRN training (GS-train in Figure 1), for fitting RNA half-lives (GS-fit in Figure 1), and for validating the predicted interactions (GS-fit or GS-validate in Figure 1). The gold standard was derived by combining binding and expression information from three major sources (Table S1). The core data was taken from the YEASTRACT repository (Teixeira et al., 2006; Monteiro et al., 2008; Abdulrehman et al., 2011; Teixeira et al., 2014), which is a curated repository of > 200,000 regulatory interactions in yeast, obtained from > 1,300 bibliographic references. The repository contains two types of evidence for each potential regulatory interaction: *direct* and indirect. Direct evidence denotes an interaction coming from an assay that directly established a physical binding event, such as ChIP-seq or one-hybrid assay. *Indirect* evidence comes from differential expression analysis after a TF knock-out or overexpression assay indirectly suggests a regulatory relationship. We first filtered these data to obtain a conservative list of 2,577 regulatory interactions that have one source of direct evidence and two sources of indirect evidence. At this stage, these interactions were unsigned, i.e. they did not include information about whether the regulatory interaction is positive or negative.

**Figure 1:**
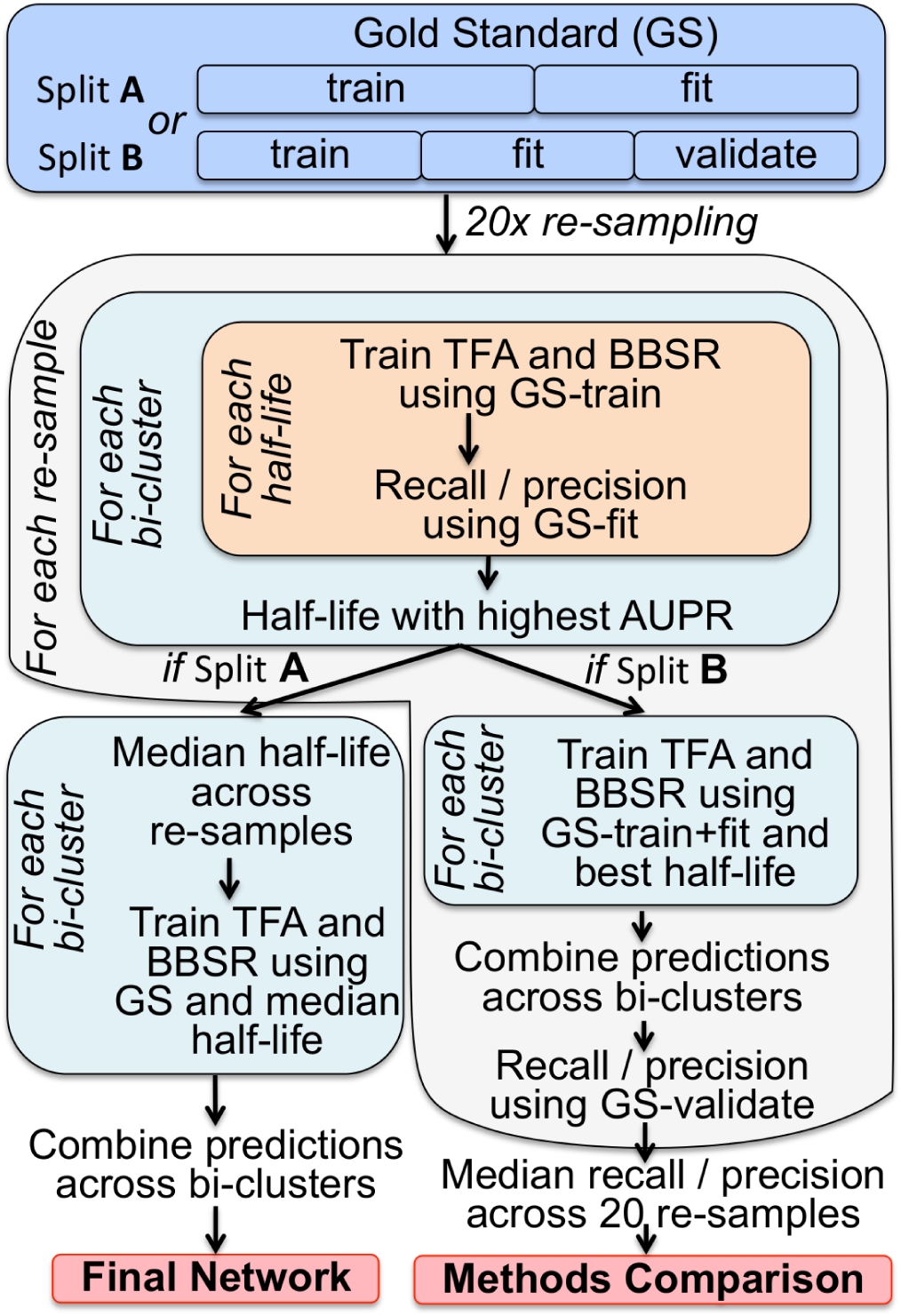
Outline of the workflows employed in this paper. We use two strategies for splitting the Gold Standard (GS) of interactions. Split **A** approach is used to make optimal predictions of condition-and gene-specific RNA half-lives. These half-lives are furthermore used to create the final predicted network. Split **B** approach is used to evaluate the improvement in network inference accuracy conferred by bi-clustering and estimating RNA half-lives. In both cases, GS was randomly re-sampled into two or three equal sets of interactions, with GS-train used to estimate Transcription Factor Activity (TFA) and run Bayesian Best Subset Regrssion (BBSR), and GS-fit to find the optimal condition-and gene-specific prediction of RNA half-lives, based on maximum area under precision-recall curve (AUPR).

As TFA estimation performs best when all prior known interactions are signed (see Section 6.1.6.1), we processed the list further to maximize the number of signed interactions. YEASTRACT provides information on the signs for some interactions, e.g. those derived from expression analysis of knock-out mutants. To add signs from the YEASTRACT database, we used the following rule: a regulatory interaction was deemed "positive" if the target gene was down-regulated upon TF knock-out, and "negative" if the opposite were the case. As some interactions were detected in multiple experiments with opposite sign annotations, we only considered the signs that were measured in assays conducted under normal conditions, labeled as "YPD medium; mid-log phase" in the YEASTRACT database. In case there was still conflict, we employed the majority rule, and in case of a tie, the interaction was discarded (set to 0). This procedure resulted in 1,155 signed interactions in total.

To expand this dataset, we obtained additional regulatory interactions from the Saccharomyces Genome Database (SGD), curated by biocurators (Cherry et al., 2012; Costanzo et al., 2014) and from the Kemmeren et al. (2014) dataset of 1,484 knock-out experiments (Kemmeren et al., 2014). These interactions were only used to assign signs to interactions that were still unsigned in the list of 2,577 interactions that had one direct and two indirect evidence types in YEASTRACT. These additions expanded our list of signed interactions to 1,403.

These 1,403 interactions constitute the set of signed prior known interactions used throughout this paper, which we denote as the *Gold Standard* (GS). Section 6.1.4, Figures S2 and S3, and Supplementary File SuppDoc1.pdf describe more details about the creation of GS and test it against other collections of interactions.

### 2.4 Inferelator Model

We used and modified code for the Inferelator version 2015.03.03 (Bonneau et al., 2006; Greenfield et al., 2013; Arrieta-Ortiz et al., 2015). We describe the Inferelator core model in this section, and more details can be found in Section 6.1.6. The Inferelator algorithm calculates the optimal model of regulation for each target gene independently of other genes. The model for each gene *i* is based on the assumption that the dynamics of transcription regulation are governed by the following relation:

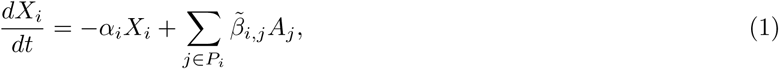

where *X_i_* is the RNA expression level of gene *i*, *P_i_* is the set of potential regulators of gene *i*, *A_j_* is the activity of TF 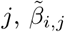 is the coefficient of regulatory interaction between TF *j* and gene *i*, and *α_i_* is the RNA degradation rate of gene *i*.

To estimate the parameters 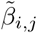 we can approximate Equation 1 using finite differences, and divide both sides by *α_i_*:

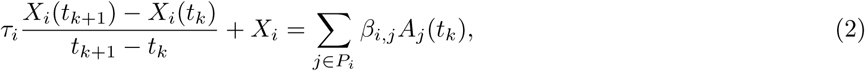

where the time axis *t* has been broken up into discrete time points at which the data was collected, indexed by *k*. The left hand side of Equation 2 is the *response variable*, whereas the the right hand side is the *design variable*. Note that 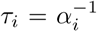, and is related to the RNA transcript half-life HL_*i*_ via HL_*i*_ = *τ*_i_ log (2), and 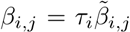. Also note that throughout our analysis, no corrections for cell division times were made, as it was impossible to determine them for each of the 2,577 experiments coming from 119 labs. Given that median half-lives are much shorter than doubling times, the effect is tolerable.

The response variable is first used together with prior known interactions to calculate TF Activities (TFA) for every TF (Section 6.1.6.1). TFA is derived from expression changes of the prior known targets of a TF, and has been shown to improve TRN inference dramatically in prokaryotes (Arrieta-Ortiz et al., 2015). The same prior known interactions are then used in a constrained regression step that selects the most likely model of regulation for every gene using a data-driven approach called *Bayesian Best Subset Regression* (BBSR). For calculating TFA and BBSR, we used our Gold Standard or a subset thereof as prior known interactions. Figure 1 and Section 2.5 outline the workflows employed in this paper, specifying how the Gold Standard was used in each of them. The final output of the Inferelator is a list of confidence scores for all possible regulatory interactions, determined using a computational knock-out assay. Each Inferelator run was performed on 50 bootstraps of the RNA expression data, and the final confidence scores for all interactions were computed by rank-combining the conffidence scores across bootstraps. For more detail, see Section 6.1.6.2.

### 2.5 Subsampling the Gold Standard for RNA-half Life Fitting and Error Estimation

To use our Gold Standard for both parameter fitting and method evaluation without overfitting, we designed two strategies for re-sampling the Gold Standard (Figure 1). For assessing the dependency of inference accuracy on RNA half-life (Figures 4 and 5A) and obtaining optimal gene-and condition-specific RNA half-lives (Figures 5B and 6, S4 and S5), we used Split A. This method involved randomly selecting a pre-speci ed fraction of Gold Standard interactions to be in the training set (GS-train), with the rest of the interactions to be used for fitting half-lives (GS-fit). We set the fraction of data used in the ‘training’ set to 0.5, although our results also hold for other values (Figure S9). This procedure was repeated 20 times, and for each re-sample, RNA half-lives were fit as described in Section 2.6.

To assess whether fitting condition-and gene-specific RNA half-lives in this manner improves performance (Figures 7A and 7C-7D, Table 1), we used Split B, where a third set of GS interactions (GS-valid) was held out and used only for estimating accuracy of our algorithm’s predictions. We created GS-valid to avoid over-fitting, and it was exclusively used to estimate the accuracy of the network computed using prior known interactions in GS-train and half-lives obtained using GS-fit. Each interaction was assigned one of the three categories randomly (GS-train, GS-fit, or GS-validate), with probabilities 0.34, 0.33, and 0.33, respectively. This way of splitting the Gold Standard was also applied to the other methods (Genie3 and iRafNet) for benchmarking purposes, keeping the random assignments of interactions into GS-train, GS-fit, and GS-valid identical across the methods for each GS re-sample.

**Table 1:**
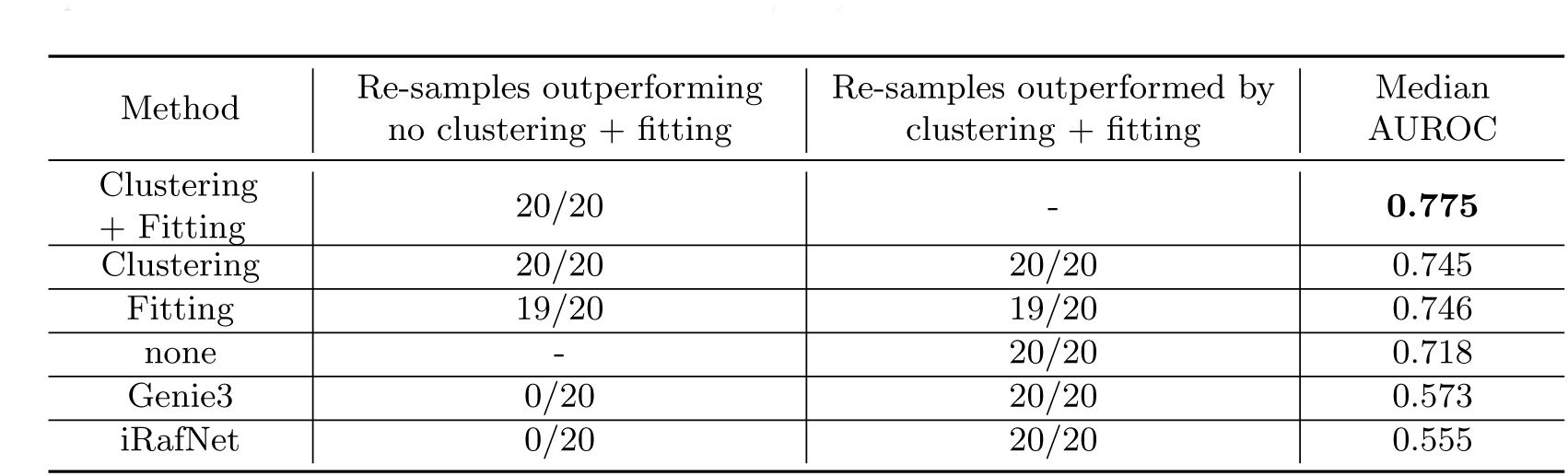
Both expression data bi-clustering and RNA half-life tting independently improve Inferelator performance (second column). Furthermore, combining the two modifications improves performance as compared to using either one of them separately (third column). Columns 2 and 3 show the number of times the AUPR measured on GS-fit from the same re-sample was higher for one method than the other, given a pair of methods specified by the row and the column. Median AUPR for each approach is reported in the fourth column. See Sections 2.5, 2.6, and 6.1.7 for further details.

We used two measures of network prediction accuracy to assess the quality of our predictions: Area Under Precision-Recall curve (AUPR) and Area Under the Receiver Operating Characteristic curve (AUROC). The two measures were calculated in the standard way, as described in Section 6.1.6.3. We focus here on AUPR, as it is more sensitive for high-scoring interactions compared to AUROC, which distributes the weights more equally across the entire list of predictions. Therefore, a model with maximal AUPR is desirable for small-scale, targeted validation experiments. AUROC is also inferior to AUPR in the class-imbalanced (skewed) regime, in which the sizes of true positives and false positives differ substantially (Davis & Goadrich, 2006), which is the case for our data.

### 2.6 RNA Half-Life Estimation

The primary advance described here is the explicit modeling and incorporation of RNA degradation rates into the large-scale inference of TRN. To do so, we first developed a procedure for comparing network prediction between models that assume different RNA half-lives (RNA half-lives are inversely proportional to RNA degradation rates, see Section 2.4).

As shown in Figure 1, Split A involves assembling two sets of interactions from the GS: one for training TFA and BBSR (GS-train), and one for calculating AUPR (GS-fit). We pre-specified values of τ that we used is τ = 0, 5, 10, 20, 30, 40, 50, 60, 70, 80, 90, 100, 120, 140, 160, 200, and 250 minutes, designed to span the range of known RNA half-lives (Neymotin et al., 2014; Munchel et al., 2011; Sun et al., 2013; Schwalb et al., 2012; Miller et al., 2011).

The Inferelator was run while setting τ to a given value on the list for every gene and every condition either in the given bi-cluster or in the entire data set, using GS-train as prior known interactions. Precision and recall curves were computed for each Inferelator run corresponding to a re-sample and a value of τ, treating GS-fit as the set of true interactions. Comparisons of precision-recall curves across RNA half-lives were made by taking the element-wise median of the precision and recall vectors across GS re-samples for a given value of RNA half-life (Figure 4A). Comparisons between AUPRs measured for different τ’s were made while keeping GS-train and GS-fit constant for each re-sample, as represented by isochromatic curves in Figures 4B and 5A, and Figure S4, and an optimal τ was chosen by maximizing AUPR. We also compared performances between models with different RNA half-lives using AUROC instead of AUPR, yielding similar optimal half-lives (Figures S6, Figures S7, Figures S8).

Finally, our RNA half-life predictions for each condition and gene bi-cluster were made by considering the distribution of optimal half-lives across the 20 re-samples for a given gene and condition bi-cluster, using the Split A procedure in Figure 1. These distributions are shown in Figure 6 for various gene and condition clusters, and their medians are shown for every bi-cluster in Figure 5B. These median values of AUPR constitute our RNA half-life predictions for each gene and condition bi-cluster. To predict RNA half-lives of translation genes (Figure 6C and S5), we separately applied the same AUPR maximization procedure to each condition cluster, using only cytoplasmic translation genes and their known regulators for AUPR calculations, because the entire gene cluster that was enriched in translation genes had too many genes that were not related to translation. This was not an issue for nucleotide metabolism genes, so the entire gene cluster was used to predict their optimal half-lives. Supplementary File SuppData3.zip contains the final RNA half-life predictions for each gene and condition cluster. For more detail, see Section 6.1.5.

To estimate the improvement in TRN inference accuracy from fitting and incorporating bi-cluster-specific RNA degradation rates, we first split the Gold Standard according to Split B, into GS-fit, GS-train and GS-validate. For a given re-sample, the predicted RNA half-lives were determined by maximizing AUPR on each bi-cluster. Using those values of RNA half-lives and the same re-sample of the Gold Standard, the Inferelator was trained again, but now using a union of GS-train and GS-fit (GS-train+fit) for TFA and BBSR computation. The final precision-recall curve as reported in Figure 7A was calculated by adding confidence scores across condition clusters for each re-sample, calculating precision and recall on the GS-valid set corresponding to that re-sample, and then taking the element-wise median of the precision and the recall vectors across the 20 re-samples. Figures 7C and 7D were calculated the same way, but with true positives and false positives instead of precision and recall in the last (validation) step. Note that the magnitude of the increase in inference accuracy due to half-life fitting that we calculate using the Split B approach (Table 1) is an underestimate of the actual increase in accuracy of our final predicted network, which is produced using the Split A approach (see Section 6.1.7 for more detail).

### 2.7 Function Enrichment Analysis

For determining Gene Ontology (GO) enrichments, we employed the hypergeometric test using the hyperGTest function from the R Bioconductor package ‘GOstats’, and the R Bioconductor package ‘org.Sc.sgd.db’ (Carlson et al., 2014). We used the 997 genes that have at least one interaction in the GS as the background for the hypergeometric test. The p-values obtained from the hypergeometric test were corrected for multiple hypothesis testing using the Bonferroni correction. For each gene cluster, we reported the top GO enrichment terms, all of which were statistically signi cant (p < 0.05). For the full list of gene ontology terms enriched in each cluster, see Supplementary File SuppData1.zip.

### 2.8 Combining Condition-Specific Networks to Create the Final Yeast Network

The final predicted network of regulatory interactions was created in several steps (Figure 1). First, bi-cluster specific RNA half-lives were determined by re-sampling the Gold standard into GS-train and GS-fit 20 times (i.e. employing the Split A approach). The predicted half-life for each bi-cluster was determined by taking the median across the half-lives that optimized AUPR in each of the 20 GS re-samples. Then the Inferelator was trained on each bi-cluster using the full Gold Standard and the predicted value of the corresponding RNA half-life. The final combined confidence scores were obtained by adding the confidence scores across the condition clusters for every gene. To convert the final (combined) ranked list of interactions into a set of predicted interactions, we set a cutoff at precision=0.5, as calculated on the full GS. This resulted in 1,462 new (i.e. not present in the GS) interactions, which can be found in Supplementary File SuppNetwork1.tsv. Supplementary File SuppNetwork2.tsv contains all predictions and their precision values.

## 3 Results

### 3.1 Assembly of Comprehensive Data Sets for High-Quality Inference

One of the main advantages of using baker’s yeast to study eukaryotic TRNs is the rich availability of existing literature that characterizes yeast regulatory phenomena in a broad sample of experimental conditions. To produce a comprehensive and accurate yeast regulatory network, we assembled all components, such as a list of 563 transcription factors (TF), the Gold Standard of interactions, and the RNA expression data set. The expression data set originated from 119 labs and spanned a wide range of experimental conditions, but used the same transcriptomics platform throughout. Comprised of 5,716 genes and 2,577 samples, as shown in Figure 2, it is one of the largest expression data sets used in network inference in yeast.

**Figure 2:**
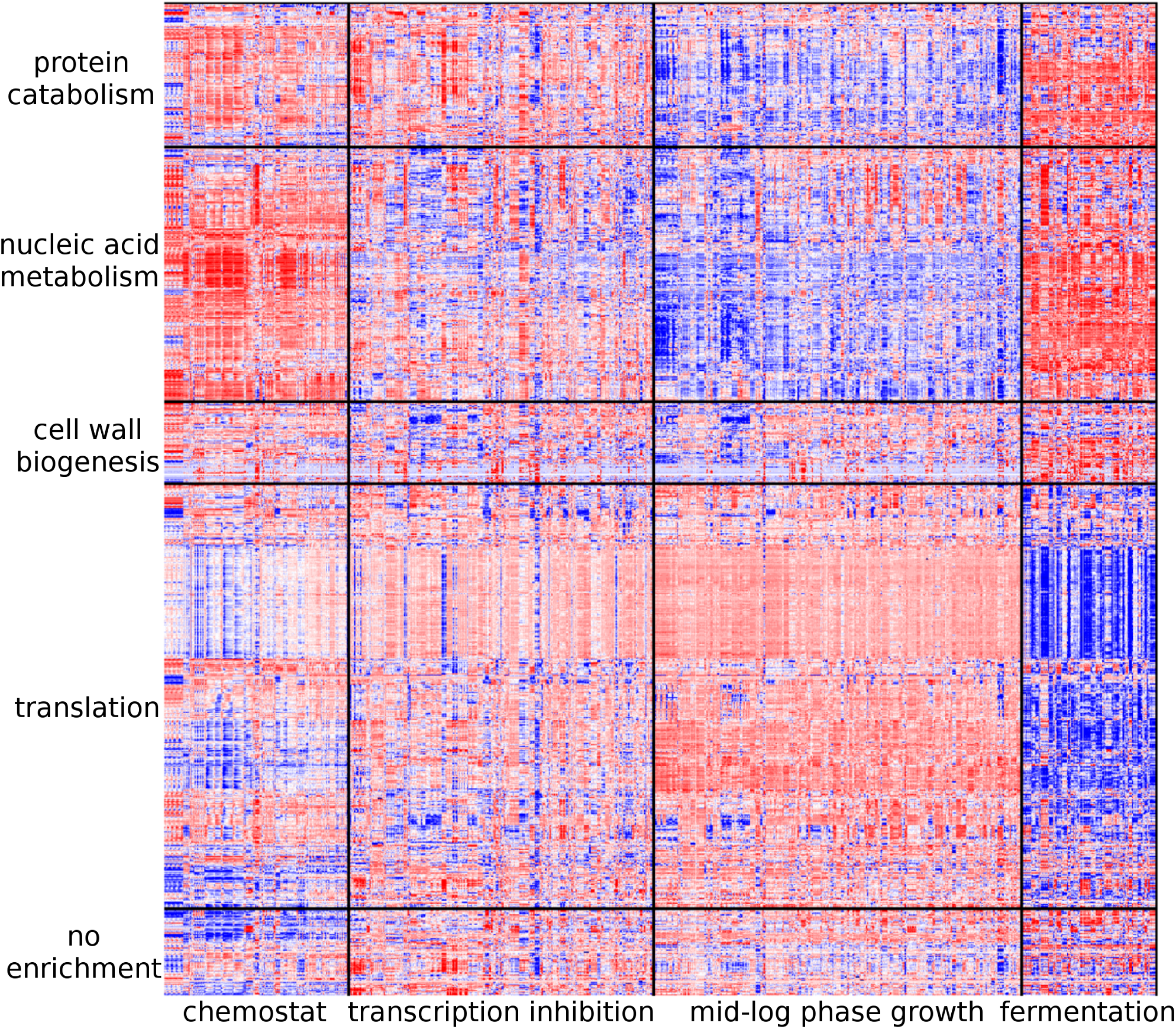
A heatmap of RNA expression data used for in this study. Genes are positioned on the vertical axis, whereas conditions are positioned on the horizontal axis. Bright red color denotes higher expression levels, whereas dark blue denotes lower expression. The four conditions clusters are shown at the bottom, whereas the five gene clusters are labeled on the left. The names of gene clusters correspond to a GO category that was most highly enriched in each cluster. Only the 997 genes that are also present in the GS are shown, although our final network was derived from expression data of 5,716 genes.

The Gold Standard (GS) of regulatory interactions combines multiple types of regulatory evidence from multiple databases (Table S1) and includes 1,403 signed interactions (i.e. distinguishing between activation and repression).

This number represents only a small fraction of all potential regulatory interactions in yeast, but each interaction is confirmed by at least three sources of evidence: one of which is direct (e.g. ChIP-chip) and two are indirect (e.g. based on TF knock-out expression changes). The GS is therefore highly enriched in true positives. Figure S2 shows that this strategy-maximizing conffidence rather than number of interactions in the GS - results in more consistent regulatory network models than other approaches.

Another development addresses the notion that several discrete modes of gene expression regulation are employed in responding to different categories of stress (Lehtinen et al., 2013; Hart et al., 2015; Yang & Leskovec, 2014). Indeed, when clustering the entire expression data set, we identified four clusters with distinct expression characteristics (Figure 2). Figure 3 shows the characteristics of each condition cluster, as extracted from the associated meta-data. Condition cluster 1 is related to chemostat growth environment under which nutrients are continuously supplied. Experiments include nutrient limitation and long time-series. Condition cluster 2 is related to transcriptional inhibition and oxidative stress experiments. Many of the transcriptional inhibition studies were early attempts to measure RNA degradation rates by inhibiting Pol II (Shalem et al., 2008, 2011). Cluster 3 is enriched in mutant-wildtype comparisons for cells grown in a rich medium. Finally, condition cluster 4 includes alcoholic fermentation experiments involving industrial strains and processes of producing wine and beer. The time series experiments in this condition cluster often last several days.

**Figure 3:**
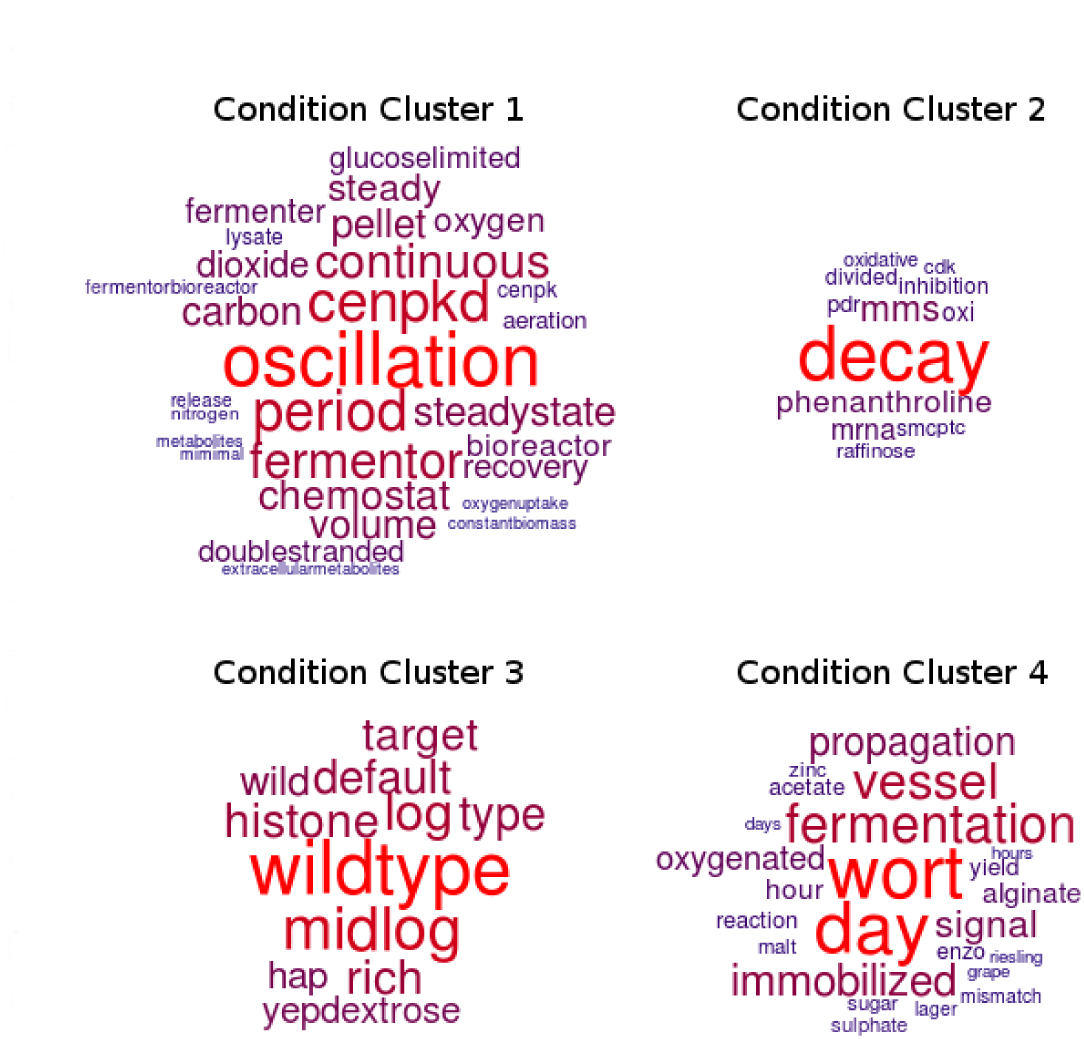
Terms enriched in each condition cluster. Words that are most highly enriched, as determined by the procedure described in the main text and in Methods, are displayed for each cluster. The higher the enrichment, the larger the font. The same cutoff of *p* < 10^−20^ (after the Bonferroni correction) was used for all four clusters.

Finally, we also clustered the data with respect to similarities between gene expression patterns, and analyzed the clusters for function enrichments. The most significant enrichments included cytoplasmic transl-ation, cell wall biogenesis, nucleic acid metabolism, and protein catabolism across four of the five clusters, respectively (p-value < 0.05, Figure 2). Combining condition and gene clustering resulted in 20 bi-clusters of the RNA expression data matrix.

### 3.2 Global and Local Network Prediction is Sensitive to RNA Stability and Recapitulates Known Trends

To assess whether the accuracy of the predicted network is sensitive to RNA degradation rates, we first ran the Inferelator on the entire unclustered expression data set for a range of preset RNA half-lives and plotted the different precision-recall curves as shown in Figure 4. We found that an RNA half-life between 20 and 50 minutes maximizes the area under the precision-recall curve (AUPR). Intriguingly, the optimal half-life range is consistent with experimental measurements, which estimate median RNA half-lives in unperturbed conditions between 10 and 15 minutes (Miller et al., 2011; Neymotin et al., 2014; Munchel et al., 2011).

**Figure 4:**
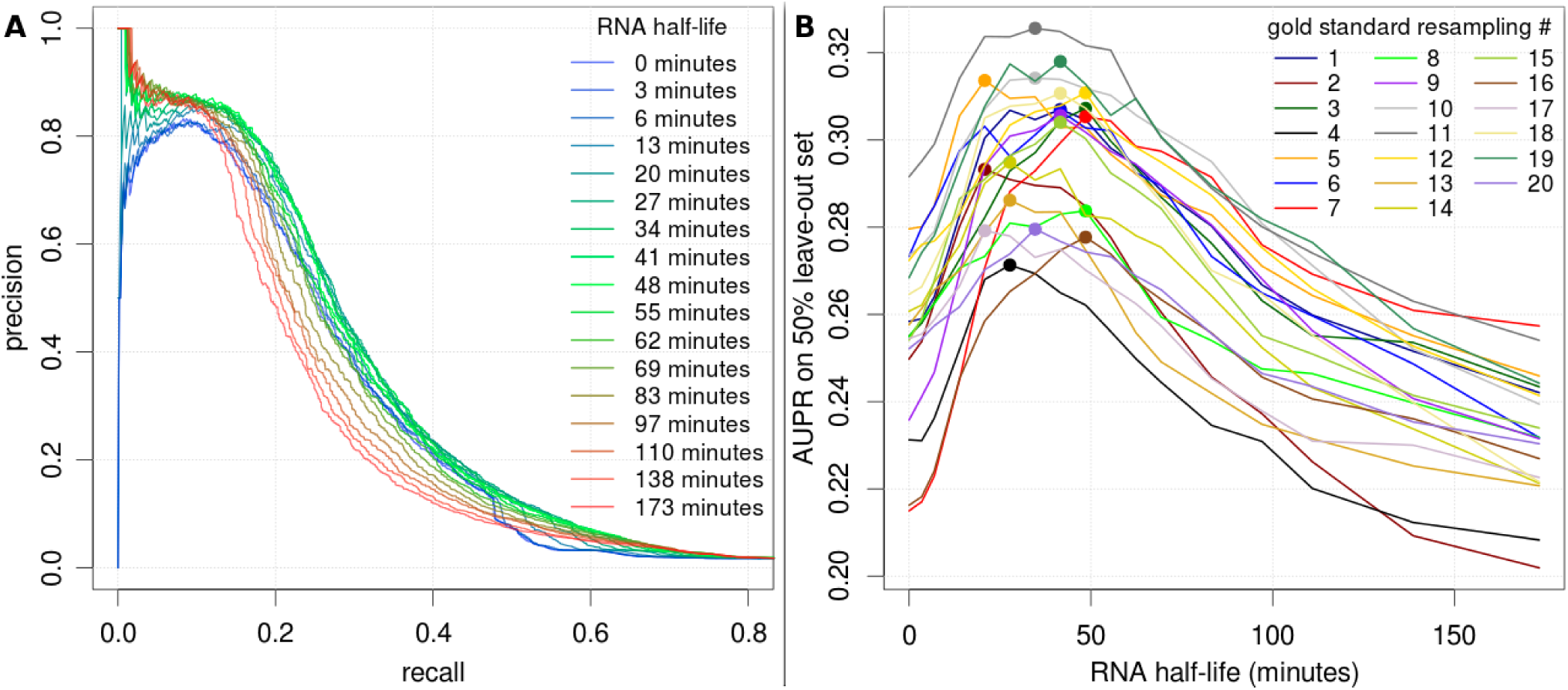
Network inference is sensitive to RNA half-lives. A) Precision-recall curves on the Inferelator output, with each line corresponding to a different pre-set value of RNA half-life. Each line displays the median precision and recall across 20 Gold Standard (GS) re-samples. B) Area under the precision-recall curve (AUPR) as a function of pre-set RNA half-life. Different lines denote 20 independent GS re-samples, and colored dots represent the maximum AUPR for a given GS re-sample.

As the data used in our work spans a variety of conditions very different from unperturbed cells growing inrich medium, we proceeded to optimize RNA half-lives for the condition clusters separately. Figure 5A shows that the optimal RNA half-life differs between condition clusters. The distribution of optimal half-lives varies in width depending on the condition cluster, suggesting a variation in half-life specificity in different conditions. The lower optimal RNA half-lives for the "chemostat" and "log-phase growth" condition clusters-which represent conditions that are perturbed less severely than the "transcription inhibition" cluster, and represent typical laboratory strains (unlike the "fermentation" cluster)-are consistent with the observed median RNA half-lives of 10 to 15 minutes in unperturbed laboratory strains.

**Figure 5:**
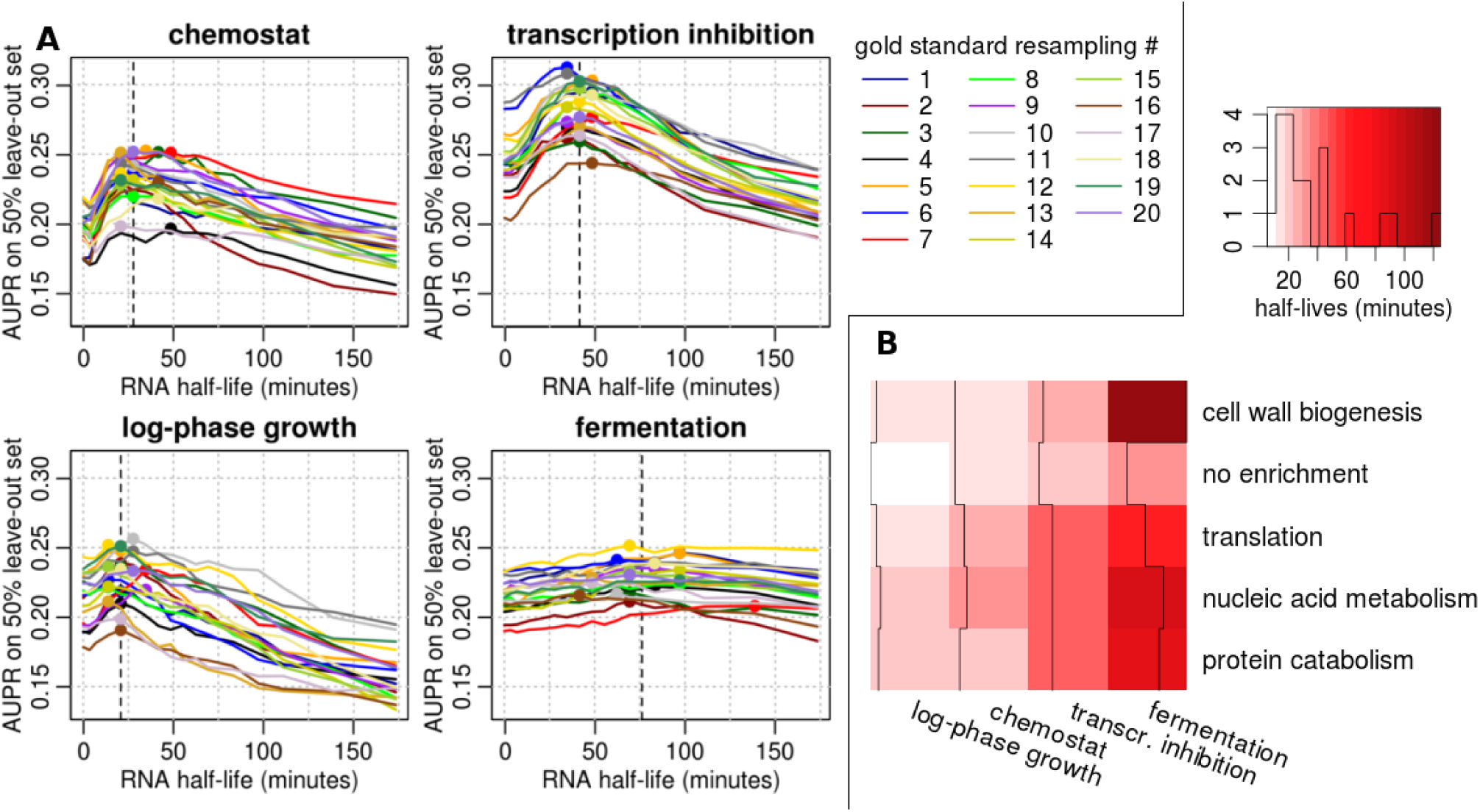
Network inference accuracy is sensitive to RNA half-lives in a condition-and gene-specific manner. A) AUPR as a function of pre-set gene-independent RNA half-life, where different colored lines correspond to 20 independent GS re-samples. The four panels correspond to condition clusters. Colored dots denote optimal half-lives, and horizontal dotted lines correspond to the median optimal half-life across re-samples. B) Optimal half-life for each condition and gene bi-cluster. Vertical black lines trace the magnitude of half-lives across gene clusters for every condition cluster. For the full plot of AUPR trajectories for every bi-cluster, see Figure S4.

To assess the dependence of inference accuracy on half-life for each gene cluster, we extended this analysis to the 20 bi-clusters. Figure 5B summarizes the bi-clustering results by showing a heatmap of median values of half-life that optimized the AUPR on that bi-cluster. Figure S4 shows that AUPR trajectories for each bi-cluster peak at distinct values of RNA half-life.

The bi-cluster specific, optimal RNA half-lives represent gene-and condition-specific RNA half-life predictions and recapitulate known biology (Figure 6). First, the range of predicted half-lives under normal conditions is similar to the range of experimentally measured half-lives reported in recent experimental measurements under normal conditions (Figures 6A and E). The most prominent pattern in Figure 5B is the increased half-lives in the “fermentation” condition cluster compared to the other condition clusters. We hypothesize that longevity (or slow growth rates) correlates with long RNA half-lives.

**Figure 6:**
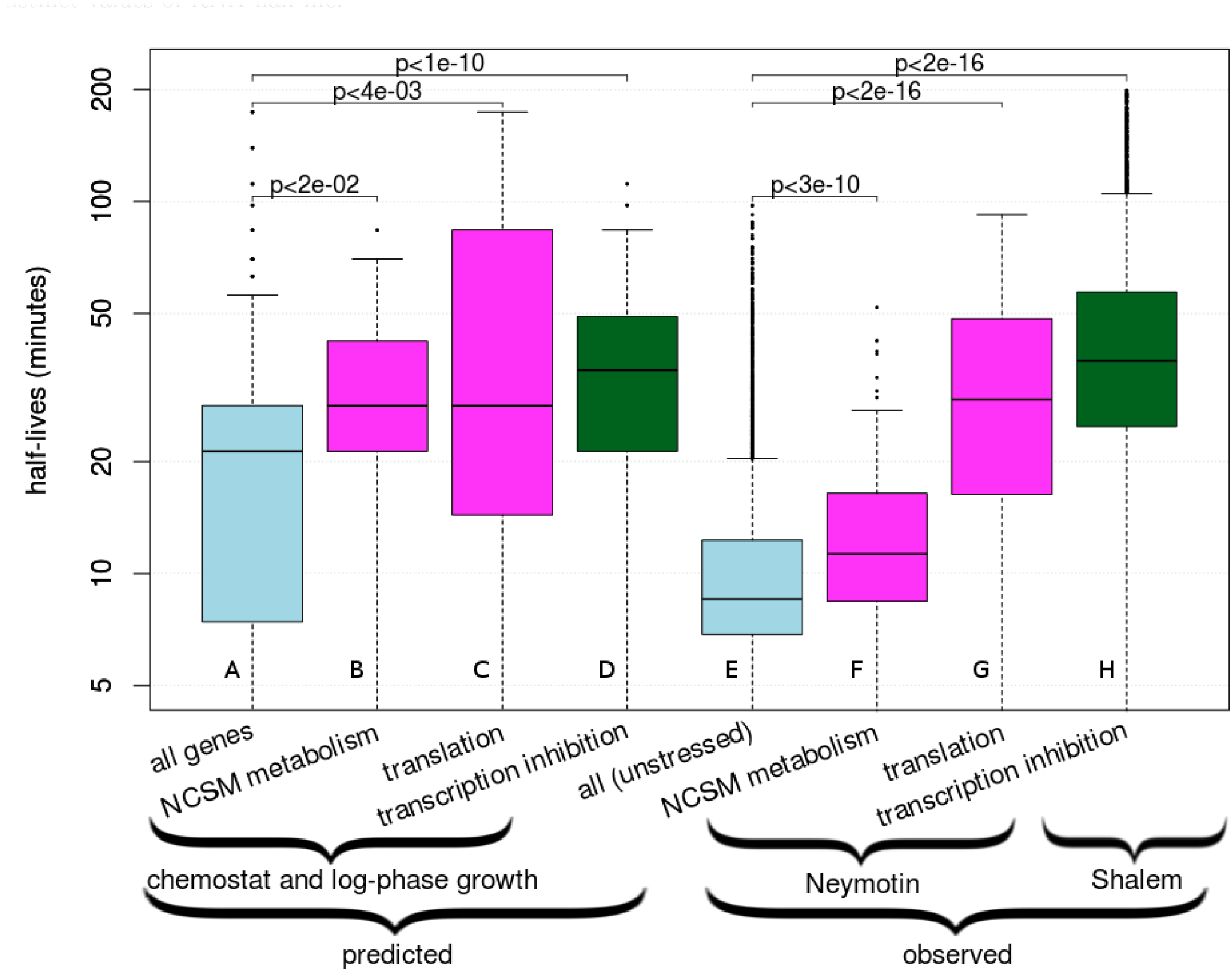
Predicted and observed differences in distributions of RNA half-lives between different groups of genes and conditions. A-D show predicted values, and E-H show experimentally measured values. Predicted values reflect distributions of RNA half-lives across the 20 GS re-samples. Magenta color denotes subsets of genes whose half-lives were predicted in non-extreme condition clusters (“chemostat” and “log-phase growth”) or measured in normal external conditions (Neymotin et al., 2014). These gene clusters, nucleotide metabolism (NCSM) and translation, are defined in the main text. Light blue denotes all genes predicted or measured in non-extreme or normal conditions, respectively. Green denotes half-lives of the entire transcriptome, predicted for the “transcription inhibition” condition cluster or measured under conditions that inhibited transcription (Shalem et al., 2008).

The next most prominent pattern in Figure 5B are the long half-lives in the “transcription inhibition” cluster compared to the “log-phase” and “chemostat” condition clusters (Figure 6D vs. Figure 6A, Wilcoxon p < 10 −^10^). This condition cluster is enriched in studies that measured RNA decay rates by inhibiting transcription. Indeed, methods that use transcription inhibition, e.g. Shalem et al. (2008), identify significantly longer RNA half-lives than those using less invasive metabolic labeling methods, e.g. Neymotin et al. (2014), as is shown in a comparison between Figure 6H and Figure 6E (Wilcoxon p < 210^16^), (Neymotin et al., 2014; Pelechano & Pérez-Ortín, 2008). This phenomenon is also closely related to buffering, wherein increased transcription rates in mutant strains correspond to decreased transcript stability, and vice versa (Sun et al., 2012).

Another example illustrating successful prediction of RNA half-lives is shown for ribosomal genes, which are known to be more stable than other genes under normal conditions (Neymotin et al., 2014; Munchel et al., 2011). Again, the prediction of RNA half-lives in our framework-which is completely independent of experimental half-life measurements and is exclusively based on expression data and network priors - confirms this bias. The predicted half-life for the 115 ribosomal genes in the "translation" gene cluster is significantly higher than that of other genes (Figure 6, Wilcoxon p < 4<10^−3^, see Section 2.6).

Finally, the most prominent pattern in the "chemostat" and "log-phase" condition clusters is the "nucleic acid metabolism" gene cluster with very high half-lives (Figure 5, 6, Wilcoxon p < 0.02). Genes in this cluster are enriched in nucleobase-containing small molecule (NCSM) metabolism. Experimentally measured half-lives of these 207 genes confirmed this trend: under normal conditions, the genes exhibited higher RNA half-lives compared to all genes (Wilcoxon p≤5<10^−10^, Figure 6F). Therefore we demonstrated that optimizing TRN inference over the biophysical RNA half-life parameter accurately predicts both known and novel condition-and gene-specific trends, confirmed by direct experimental measurements.

### 3.3 Global Network Inference is Improved through the Use of Optimized RNA Stability

Having shown that the RNA half-lives optimized in our prediction are biologically relevant, we proceeded to use these half-lives to enhance prediction of regulatory interactions. To do so, we used separate, randomly chosen subsets of the Gold Standard to train the model, fit RNA half-lives for individual bi-clusters, and validate the predictions (Figure 1, Split B). A separate comparison was made for each of the 20 GS re-samples across each pair of competing methods (Figure 7, Table 1). These comparisons were made in terms of AUPR, as calculated on the respective GS-valid set of interactions, using the single value of half-life optimized on the respective GS-fit set of interactions. See Section 2.6 for more details on the methodology of methods comparisons.

**Figure 7:**
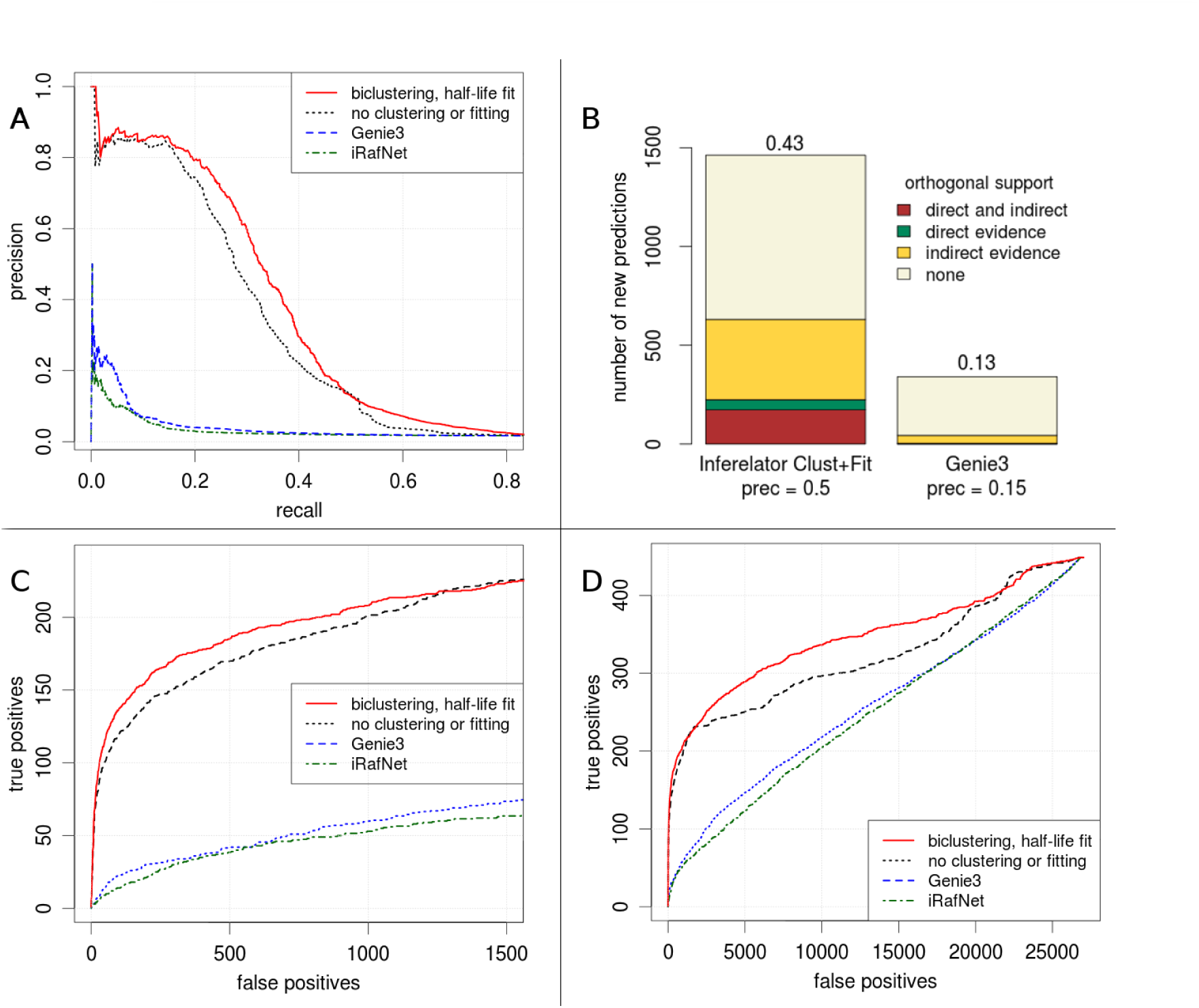
Inferring bi-cluster specific RNA half-lives improves network inference accuracy, resulting in better network predictions than other state-of-the-art methods. A) The estimated improvement in the precision-recall curve as a result of bi-clustering expression data and optimizing the half-life for each bi-cluster (red line), compared to using the Inferelator without bi-clustering or half-life optimization (black dotted line), or compared to Genie3 and iRafNet (blue line and green line, respectively). B) Number of new predicted interactions (i.e. interactions not in the Gold Standard), obtained using the optimized bi-cluster-specific half-lives and the full GS for training, compared to new predictions from Genie3. The vertical length of each color section within each bar corresponds to the number of new interactions that were confirmed by the corresponding type of evidence in an orthogonal data source. Direct evidence refers to physical binding, and indirect refers to knock-out and overexpression assays. The number above each bar denotes the fraction of new interactions supported by at least one orthogonal source. C and D show the results of the same comparison as in A by plotting false positive vs true positive rates instead of precision and recall. C) is a zoomed-in version of D.

Inference accuracy is improved drastically by incorporating bi-cluster-specific RNA half-life predictions, compared to inference without clustering of the expression data and use of optimized half-lives (Figure 7). A significant but smaller improvement is seen with each of the two modifications taken separately: either with bi-clustered expression data but without half-life optimization, or with half-life optimization but without expression clustering (Tables 1 and S2). A total of 113 of 120 pairwise comparisons provided larger AUPR for inference with these modifocations (Table 1), and the median improvement of using both modifications is ≈ 14%. Using AUROC yielded a similar outcome (Table S2). The final AUPR value of 0.33 represents an almost eight-fold increase compared to Genie3 (Table 1), an inference method that performed best in a recent network inference competition (Marbach et al., 2012). It represents a 10-fold increase compared to another method, iRafNet (Table 1) (Petralia et al., 2015). See Sections 2.5, 2.6, and 6.1.7 for further details.

To maximize the size and accuracy of the final, integrated network, we repeated the whole procedure, but used the entire Gold Standard for training, setting the half-lives for each condition and gene cluster to the predictions made using the Split A approach from Figure 1. At 50% precision, this final transcription regulatory network contained 1,462 newly predicted interactions that were not present in the Gold Standard. Of these 1,462 interactions, 631 (43%) were validated by external data that was not included in this work, derived either from direct binding evidence (Yeastract-Direct), indirect evidence (Yeastract-Indirect, SGD, Kemmeren), or both (Figure 7B). This high fraction of independently confirmed interactions suggests that the remaining 831 new interactions are also strongly enriched in true positives.

### 3.4 The High-Quality Final Network Produces Functionally Relevant Predictions

To illustrate the value of newly predicted regulatory interactions, Table S3 lists the top ten most confidently predicted targets of the ten most connected and medium connected TFs, respectively. Many of these interactions are validated by literature. Table 2 and Table S3 display interactions that are not already in our Gold Standard, but many of these interactions are present in orthogonal collections of interactions (non-bolded targets in Table 2, superscripts in Table S3). For example, SFP1 is a well-known regulator of ribosome biogenesis (Marion et al., 2004; Cipollina et al., 2008; Reja et al., 2015), and indeed, all of its top predicted targets are ribosomal subunits. Similarly, RPN4 regulates proteasome expression (Karpov et al., 2008a,b), and proteasomal genes form the majority of its most confidently predicted targets.

**Table 2:**
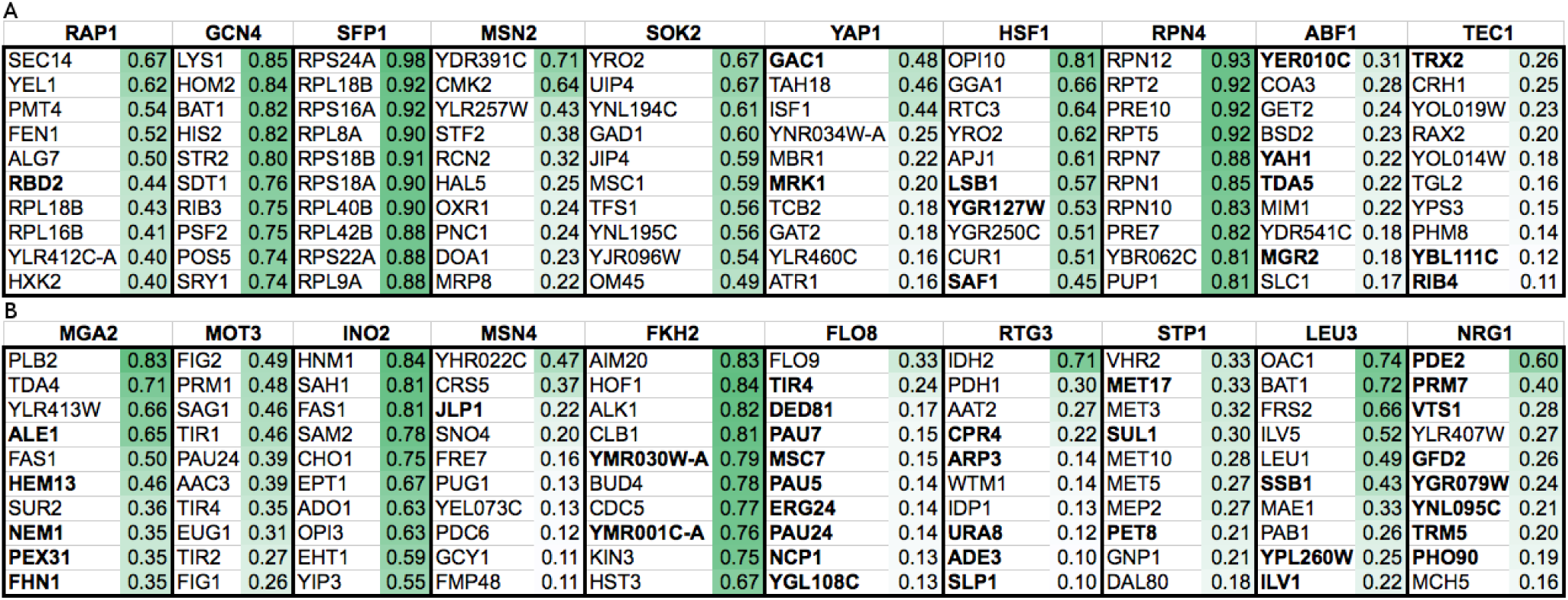
New final predictions and their precision values. A) New targets (i.e. interactions not in the Gold Standard) of the 10 TFs that have the highest number of known targets in the Gold Standard, and the precision values of these new interactions. TFs are listed in the top row, and their new targets with the corresponding precision values are listed below the TF. B) New targets of the median 10 TFs in terms of the number of known targets, out of the 97 TFs with at least one target in the GS. Precision values are calculated using the entire matrix of prediction confidence scores, containing 5,716 genes and 557 TFs. The list of true positives was defined as the entire Gold Standard. Shade of green denotes the preicision value. Bold targets correspond to interactions that were not found in any of the four major sources of interactions used in this study listed in Table S2. See Table S3 and Supplementary File SuppNetwork3.tsv for more details.

Of the 100 new interactions predicted as the top ten most likely targets of the top ten most highly-connected genes, only 13 (13%) have not been validated by one of the four regulatory interactions data bases that we employed. However, existing literature suggests that many of these completely new interactions are likely to be correct. For example, HSF1 is a key regulator of diverse stresses and monitors translation status through interaction with the Ribosome Quality Control complex (RQC) (Brandman et al., 2012). We make a completely new prediction that HSF1 regulates LSB1 and SAF1. SAF1 has four other transcription regulators (BUR6, MED6, SPT10, SUA7) all detected under various stresses, especially heat shock (Mendiratta et al., 2006; Venters et al., 2011)-suggesting that HSF1 may be a true regulator. LSB1 regulates actin assembly and prion modulation in yeast (Ali et al., 2014), but has not been previously linked to HSF1. However, several recent studies have linked HSF1 to actin assembly: one study identified altered actin cytoskeletal structures in yeast deficient in the RQC-Hsf1 regulatory system (Yang et al., 2016); another study showed that overexpressing HSF1 in worms increases actin cytoskeleton integrity and lifespan (Baird et al., 2014); a third study in mammalian cells confirmed that active HSF1 affects the actin cytoskeleton (Toma-Jonik et al., 2015). Therefore, it is tempting to hypothesize that actin assembly regulator Lsb1 is the missing link by which HSF1 a ects the actin skeleton.

Another validation for newly predicted regulatory interactions arises from gene knockout phenotypes. For example, the TEC1 transcription factor regulates filamentation genes, but also positively affects lifespan (Mösch & Fink, 1997; Garay et al., 2014). Its newly predicted target TRX2 is a thioredoxin isoenzyme involved in the oxidative stress response (Garrido & Grant, 2002; Greetham et al., 2010). A screen by Postma et al. (2009) demonstrated increase lifespan in a TRX2 null mutant strain Postma et al. (2009), again providing supporting evidence for the regulatory interaction between TEC1 and TRX2 that was newly predicted in our work.

Finally, similar reasoning supports another example: MRK1 is a newly predicted target for the YAP1 transcription factor, which is a basic leucine zipper required for tolerance to oxidative stress and cadmium exposure (Kuge & Jones, 1994; Wemmie et al., 1994; Lee et al., 1999). MRK1 is a Glycogen synthase kinase 3 (GSK-3) homolog, which activates Msn2p dependent transcription of stress response genes (Hardy et al., 1995; Hirata et al., 2003). Interestingly, a genome-wide screen showed increased cadmium levels for the MRK1 knockout, indicating that the gene is indeed linked to a YAP1 function (Yu et al., 2012). Cadmium exposure is an example of a rare environmental condition for which binding or perturbation experiments would likely be missing, highlighting the advantage of network inference from large-scale expression data of diverse experimental design.

## 4 Discussion

Genome-wide inference of transcription regulatory networks in eukaryotes is challenging. Here, we present conceptual advances over existing work (Greenfield et al., 2013; Arrieta-Ortiz et al., 2015), demonstrating, for the first time, that using biophysically relevant models that incorporate, for example, RNA degradation, improves automatic large-scale network prediction. In addition, our approach includes other substantial improvements, such as the use of a high-quality Gold Standard of regulatory interactions, and the construction of different network models across subsets of genes and conditions. Using these advances, we present a genome-wide regulatory network, which at 50% precision predicts > 1,400 new interactions, 43% of which are validated by independent data sets, and 57% are entirely new (Figure 7B).

The high-quality Gold Standard dataset of regulatory interactions is built on several benchmark datasets of regulator-target interactions (Teixeira et al., 2006; Monteiro et al., 2008; Abdulrehman et al., 2011; Teixeira et al., 2014; Cherry et al., 2012; Costanzo et al., 2014; Kemmeren et al., 2014), but improves on these sets by adding signs and accounting for confidence measures. In constructing this Gold Standard, we found that the quality, but not necessarily the size of a benchmark set of interactions improves predictions (Figure S2).

With this signed Gold Standard, we combined network predictions from several disjoint expression sub-spaces. Combining networks that were modeled assuming distinct regulatory regimes resulted in higher recovery of known interactions than when assuming the same regulatory regime across all conditions (Figure 7, Table 1). This result is consistent with the findings that both RNA degradation rates and regulatory networks experience fine-tuned and global changes in response to changing environmental conditions (Munchel et al., 2011; Miller et al., 2011; Lehtinen et al., 2013; Hart et al., 2015; Yang & Leskovec, 2014).

Most importantly, we showed that including RNA degradation in the mathematical modeling of RNA expression changes substantially boosted inference of the transcriptional regulatory network. To the best of our knowledge, the inference framework that we used, the Inferelator, is the only approach capable of doing so on a genome-wide scale. Notably, we learned RNA degradation rates directly from time-series and steady-state expression data, without the use of values known a priori. The resulting optimal rates are surprisingly similar to experimentally measured rates and accurately reflect known trends. For example, ribosomal RNAs are more stable than other gene transcripts under normal conditions, and transcription inhibition appears to correlate with degradation globally, as has been shown experimentally (Neymotin et al., 2014; Sun et al., 2012; Pelechano & Pérez-Ortín, 2008) (Figure 6).

Given that it is still highly challenging to measure RNA degradation in living cells and relevant conditions, only a few such datasets exist (Miller et al., 2011; Schwalb et al., 2012; Sun et al., 2012; Neymotin et al., 2014; Munchel et al., 2011), and our framework can be used to predict missing rates. It can be used to reveal trends that have so far gone unnoticed, such as the long half-lives of nucleic acid metabolism genes (Figure 6). While the majority of predicted interactions were validated by external large-scale regulatory interaction data sets, we also illustrated that even the interactions not seen in any previous literature are biologically meaningful and supported by orthogonal evidence – e.g. YAP1 regulating MRK1, TEC1 regulating TRX2, and HSF1 regulating LRB1.

In a broader context, this work illustrates the promising prospect of more biophysically motivated modeling approaches to network inference. Other popular methods currently use techniques such as Random Forest (Huynh-Thu et al., 2010; Petralia et al., 2015), Mutual Information and Related Transfer Entropy (Margolin et al., 2006a,b), correlation (Butte & Kohane, 2000), and others. These methods do not provide a clear way to incorporate or recapitulate measurements of biophysical dynamics parameters that govern transcription regulation. The Inferelator’s explicit modeling of the RNA synthesis and degradation processes via a differential equation highlights the advantage of using this framework for TRN inference.

The results of this study encourage further developments of the Inferelator algorithm that would allow for an efficient incorporation and recovery of biophysical parameters, such as RNA decay rates and interaction terms between co-regulating TFs, a more careful separation of the transcription term into transcriptional activation and repression, which has only been done on the small scale (Noman & Iba, 2005; Liu & Wang, 2008; Bonneau & Aijo, 2016; Intosalmi et al., 2016), and modeling functional modifications of TFs that can affect their transcriptional activity. Given the growing body of literature on RNA-binding proteins (RBPs) (Hogan et al., 2008; Mittal et al., 2009; Janga & Mittal, 2011), our results also inspire potential approaches to model the RNA decay term explicitly as a sum of contributions from RNA degradation factors. Therefore, it is time to move inference of transcription regulatory networks to biophysically relevant models, and the work presented here provides an important step towards this goal.

## 6 Supplement

### 6.1 Supplementary Methods

#### 6.1.1 Primary Data Processing

The meta data for every whole-genome expression sample (GSM) was downloaded using the R function getGEO from the GEOquery package (Edgar et al., 2002; Barrett et al., 2013). These GSMs were filtered such that the final list only contained samples measured in Saccharomyces cerevisiae using the Yeast Affymetrix 2.0 platform (GPL2529). For each expression series (GSE) that contained at least one of those GSMs, a TAR file was downloaded from the GEO website on March 23, 2015, using the R function download.file. The raw CEL files for every sample in that GSE (filtering for S. cerevisiae and GPL2529) were furthermore extracted from the associated TAR file. For processing and normalizing the raw CEL files, we used the R packages affy (Gautier et al., 2004) and gcrma (Wu et al., 2004), using the functions ReadAffy and gcrma. All samples were processed simultaneously as one batch. For time series meta data, all time differences were converted into minutes. To test for batch effects, we used ComBat (Johnson et al., 2007; Leek et al., 2012), treating either laboratory of origin or experiment series as batches, but did not observe an improvement in network prediction (data not shown). All computations were made in R programming language, version 3.1.2 (R Core Team, 2014).

#### 6.1.2 Condition Clustering

Principal Component Analysis (PCA) of the entire expression matrix was performed using the R command prcomp, treating each expression sample as a 5; 716-dimensional vector. K-means clustering of the dimensionally-reduced (post-PCA) expression samples was performed using the R function kmeans, using the default parameters with nstart=25, and iter.max=1000, corresponding to 25 initial random configurations (among which the best one is automatically picked by the built-in algorithm in R) and a maximum of 1000 iterations. PCA and k-means clustering was performed on scaled expression data (mean shifted to 0 and variance scaled to 1), and all subsequent analysis was done on the resulting clusters using the original (processed and normalized, but not scaled) expression data. We also performed some of the downstream analysis using clusters that were obtained the same way but without scaling, which did not lead to any significant differences in terms of inference performance (not shown).

**Figure S1:**
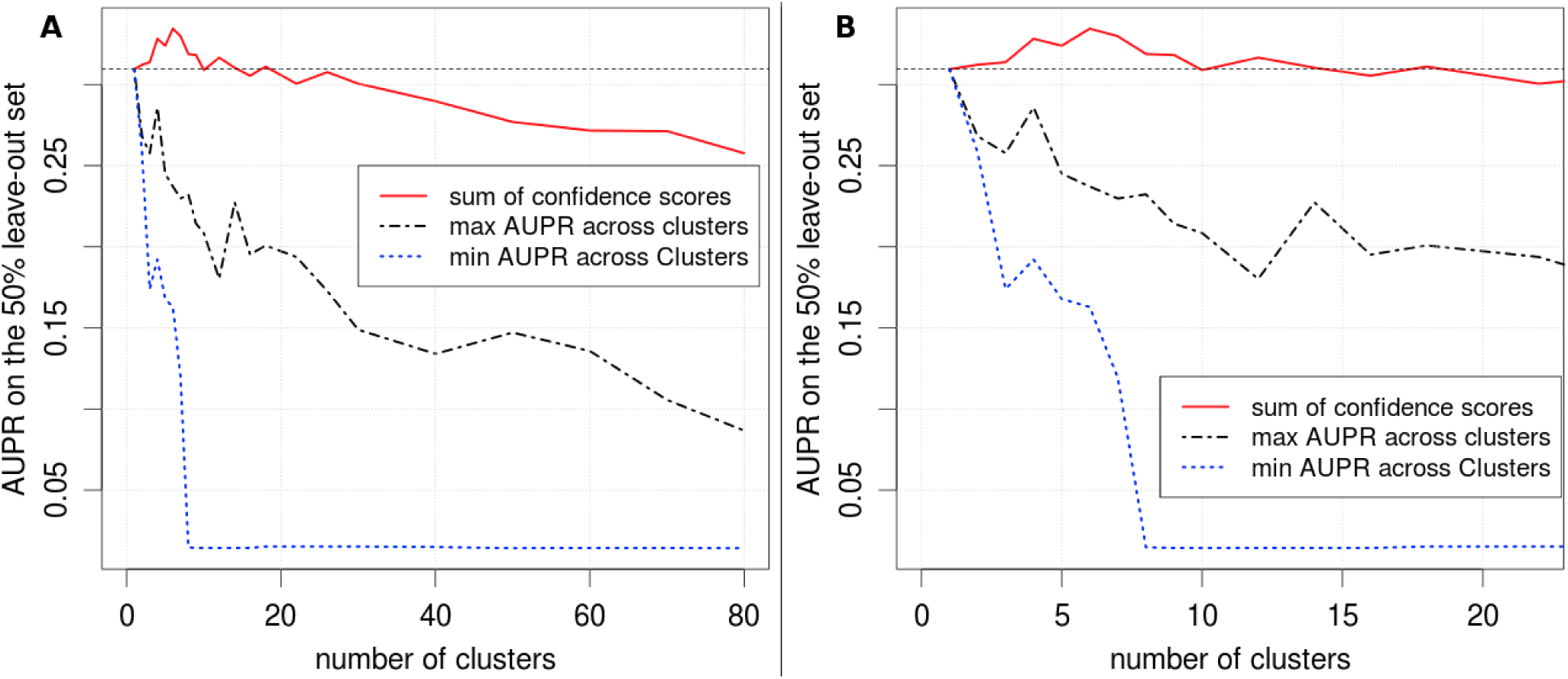
Network inference accuracy is improved when the expression data is clustered sample-wise and the predictions from the resulting clusters are combined. A) AUPR on the leave-out set is a function of the number of clusters, into which the expression data is split. B) same as A, but the x-axis is zoomed in around where the AUPR of the combined confidence scores is peaked. The red line shows the AUPR of the confidence scores combined across the clusters, the black dash-dotted line shows the highest AUPR across the clusters, and the dotted blue line shows the lowest AUPR across the clusters.

The number of condition clusters was determined by rank-combining the confidence score outputs of the Inferelator across a pre-specified number of clusters, and maximizing the prediction accuracy of this procedure as a function of the number of condition clusters. Rank-combining across the predictions obtained from a set of disjoint expression clusters was done by taking the sum of the confidence score matrices (combinedconf*.RData files). We used the AUPR of the rank-combined prediction as a proxy for prediction accuracy. The pre-specified numbers of clusters that we tested were n = 2-10, 12, 14, 16, 18, 22, 26, 30, 40, 50, 60, 70, and 80. For a given n, the entire expression data set was first clustered into n clusters (as described in the paragraph above). For each n, we ran the Inferelator separately for each of n expression clusters, using 50% of the GS as the leave-in set for TFA and model selection steps, and the rest as the leave-out set for calculating AUPR. We did not resample the GS for this analysis, we used 10 response matrix bootstraps, and set τ = 60 for this analysis. The same randomly sampled leave-in set was used across the Inferelator runs for all clusters.

Figure S1 shows AUPR as a function of the number of clusters n. Furthermore, it shows the AUPR of the worst-performing and the best-performing cluster among the n clusters. We notice that the AUPR is maximized around n = 4 and n = 7, where it is approximately equal to 0.35. We selected n = 4 as our optimal number of clusters because after n = 4, there is a sharp drop in the AUPR of the worst-performing cluster, rendering it uninformative. All downstream analysis was performed on these four clusters.

To annotate the condition clusters, all text that was provided in every available sample of the meta data table was sorted into four bins according to the sample’s assigned cluster. The following processing was done for the list of words in every cluster. First, common English words, commas, and numbers were removed. Text was converted to lower case. These operations were done using the R package tm (Feinerer & Hornik, 2015; Feinerer et al., 2008). Unnecessary whitespaces were removed, and term frequency statistics were calculated using the R package SnowballC (Bouchet-Valat, 2014). Multiple hypothesis testing for enriched terms in each cluster was performed using the Bonferroni correction. To foster visualization and interpretability, we removed spurious terms, which we did not consider biologically relevant, from these lists. The final word clouds were made using the R function wordcloud from the package wordcloud (Fellows, 2012).

In order to assign each cluster with a final label, we compared the occurrence of enriched terms in each cluster and assessed whether they come only from one lab or from multiple labs, in order to ensure that each topic is not enriched in its cluster as a result of lab bias in term usage (Supplementary File SuppData2.zip). Our condition cluster label assignments were also supported by GO enrichments in genes that had the highest relative variance compared to other genes in their cluster (not shown).

#### 6.1.3 Gene Clustering

Gene clustering was performed as described in Section 2.2. The reason for scaling expression data before clustering is that this approach results in more evenly-sized gene clusters. We performed hierarchical clustering using the R function hclust with default arguments (e.g. euclidean distance metric). We determined the number of clusters by comparing several qualities of the resulting gene clusters as a function of the number of clusters. Given a pre-specified number of clusters k, we used the R function cutree to cut the similarity tree into k clusters. For each of the k resulting gene clusters, we performed Gene Ontology (GO) enrichment analysis (see Section 2.7). After performing this analysis for k ranging from 2 to 8, we determined that setting k > 5 does not result in any new clusters with meaningful new GO enrichments as compared to the enrichments obtained with k = 5 clusters. Rather, setting k > 5 creates new clusters with no GO enrichments.

Since our method for determining gene-and condition-cluster specific RNA half-lives relies on the availability of prior known interactions with the target genes, we first performed gene clustering for the 997 genes that were present in our gold standard of interactions. For this reason, setting the number of clusters higher than k > 5 resulted in clusters with too few genes to result in significant GO enrichments. For example, with k = 5, the smallest cluster consists of 79 genes and the largest consists of 409 genes. With k = 6, the former smallest cluster was further broken down into two clusters with 44 and 35 genes in each. Small gene clusters were also unfavorable because our RNA half-life inference for a given gene cluster relies on AUPR maximization calculated on a leave-out list of prior known interactions involving the genes in that cluster. AUPR calculation gets increasingly noisy as the number of known true positives is reduced. For k = 5, the smallest number of prior known interactions with the genes in one of the five clusters was 101, and the largest was 592. This range was sufficient for producing relatively smooth half-life vs. AUPR curves (Figure S4).

For the final yeast network, we included regulatory network predictions for all 5,716 genes in our expression data set. This required us to assign each gene with one of the clusters that was determined using the 997 genes that had prior known interactions. Denote the set of those 997 genes as gGS. For a given gene g, its cluster membership was assigned by finding the gene g_prior_ 2 gGS that minimizes the euclidean distance between g and g_prior_, and assigning the cluster membership of gene g_prior_ to gene g. This approach was undertaken instead of clustering all 5,716 genes and then subsetting them to the 997 genes in *gGS* because the latter approach resulted in highly unevenly distributed cluster membership numbers among those 997 genes, which would result in noisy AUPR calculations on clustered data.

**Table S1:**
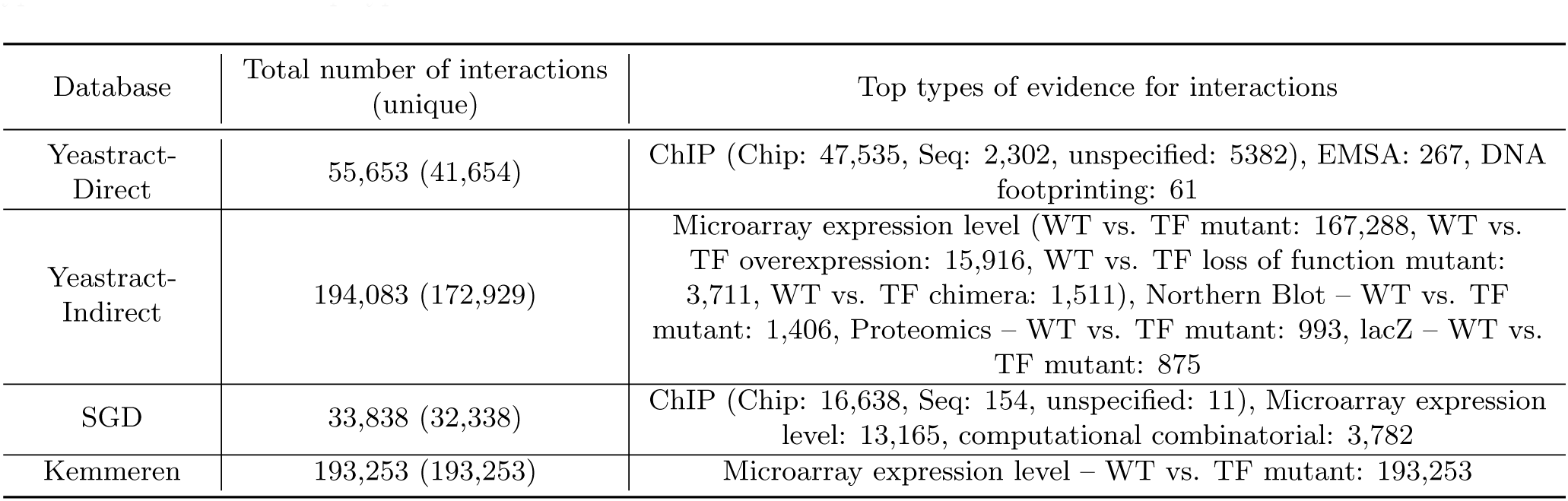
A survey of the databases of regulatory interactions used in this study. The first column denotes the name of the interactions database. The second column lists the number of interactions in each database. An interaction may be duplicated within a database if the database contains multiple entries for the same interaction (e.g. coming from different labs or different conditions), so in the parentheses we specify the number of unique interactions for each database. The third column lists the most common types of regulatory evidence in each database, including the number of interactions for each type of evidence. These top types of evidence account for over 98% of interactions in each database.

#### 6.1.4 Gold Standard Curation

Gold Stanard was created as described in Section 2.3. Here we specify some further details. In the YEASTRACT dataset, in case the sign of an interaction was specified, it appeared in the “Association Type” column. When “Association Type” had a label “Negative”, it denoted a decrease in abundance after a TF knock-out (and a positive sign according to our convention), and “Positive” denoted an increase (resulting in a negative sign according to our convention).

To obtain interactions from Saccharomyces Genome Database, we downloaded 32,338 unique regulatory interactions using Yeastmine (Balakrishnan et al., 2012; Costanzo et al., 2014). We refer to these 32,338 interactions as the SGD standard (such as in Table S3). The SGD website provided signs for 10,022 of those interactions. We observed 363 agreements and 12 disagreements in signs of the overlapping signed interactions between the list of 1,155 high-confidence signed interactions obtained from YEASTRACT (as described in Section 2.3) and the 10,022 from SGD. We used the signed portion of the SGD standard to expand our list of signed GS interactions in the following way. If the signed interaction from SGD was already in the list of 1,155 from YEASTRACT, we kept the sign that we obtained from YEASTRACT regardless of its sign in the SGD standard. If it was among the 2,577 high-confidence interactions obtained from YEASTRACT, but without a sign from YEASTRACT, we assigned it with the sign from SGD. This procedure provided an additional 117 signed interactions towards the final gold standard of interactions.

Furthermore, we used the data from Kemmeren et al. (2014), which contained the results of a study that measured genome-wide expression level changes for 1,484 knock-outs, including most TFs in yeast. First, we created a collection of interactions by considering all knock-out-induced expression changes that were reported in the original publication with a p-value of p < 0.01 after the Bonferroni correction, yielding > 193,000 interactions (which we furthermore denote as the Kemmeren standard). Of these, 131 were present but so far unsigned amongst the 2,577 high-con dence interactions described above, therefore expanding the set of high-confidence signed interactions to 1,403.

Figure S2 compares the performance of the Inferelator on the 50% leave-out set of seven different collections of interactions, when trained on the 50% leave-in set of the same seven collections (respectively). This provides a measure of consistency of a collection of interactions, estimating how effectively the Inferelator trained on one half of a collection of interactions can recover the other half of the same collection of interactions. According to both the precision-recall curve and the Receiver Operating Characteristic curve, the final Gold Standard of 1,403 signed interactions is by far the most consistent (has the highest area under the curve) as compared to any of the sources of interactions shown in Table S1. The biggest improvement in AUPR is achieved by restricting the collection of interactions from YEASTRACT to the 2,577 unsigned interactions that have 1 piece of evidence in Yeastract-Direct and 2 pieces of evidence in Yeastract-Indirect (blue line, AUPR=0.20, AUROC=0.70). Reducing those interactions to only those, for which we have evidence of the direction of the regulation in either of the three sources of interactions (YEASTRACT, SGD, and Kemmeren), further improves performance (Gold Standard, red line, AUPR=0.30, AUROC=0.73). Our Gold Standard also outperformed the MacIsaac regulatory interactions database (MacIsaac et al., 2006), which is commonly employed for evaluating network inference algorithms in yeast (Marbach et al., 2012; Siahpirani & Roy, 2016). We extended this comparison to other possible combinations of these collections of interactions (Figure S3). Our results led us to conclude that the 1,403 interactions of our Gold Standard constitute the most consistent and reliable collection of interactions in yeast known to date.

**Figure S2:**
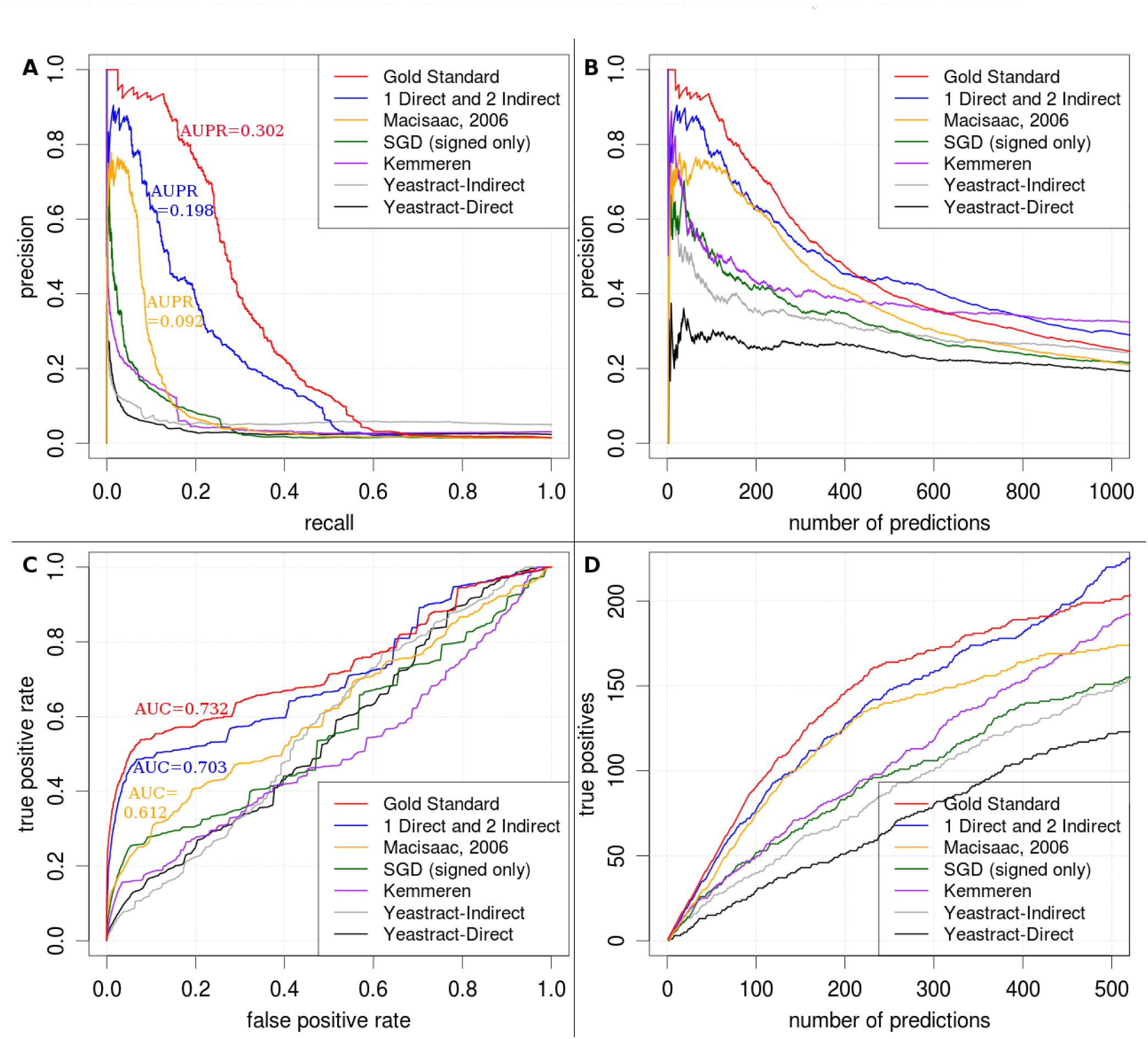
Combining multiple sources of evidence creates more consistent collections of regulatory interactions. Each of the collections of interactions listed in the legend was split into two equal parts, one for training the Inferelator and one for validation. Each line represents statistics calculated on the 50% leave-out set of the corresponding collection of interactions. Black, gray, purple, and green lines correspond to the collections of interactions described in Table S1. The green line only includes the signed portion of the SGD collection (where positive and negative regulation is pre-specified). Blue line corresponds to the interactions that appear at least once in the Yeastract-Direct collection and at least twice in the Yeastract-Indirect collection, without positive and negative regulation specified. The red line shows the final signed Gold Standard we use throughout this study. A) Precision vs. recall for the six collections of interactions. B) Precision as a function of number of predictions. C) Receiver Operating Characteristic curve for each collection of interactions. D) Number of true positives as a function of the number of predictions. Note that in B and D, performance on the Gold Standard decreases more quickly after several hundred interactions because the positives exhaust all true interactions more quickly due to the smaller size of the Gold Standard as compared to other collections shown.

**Figure S3:**
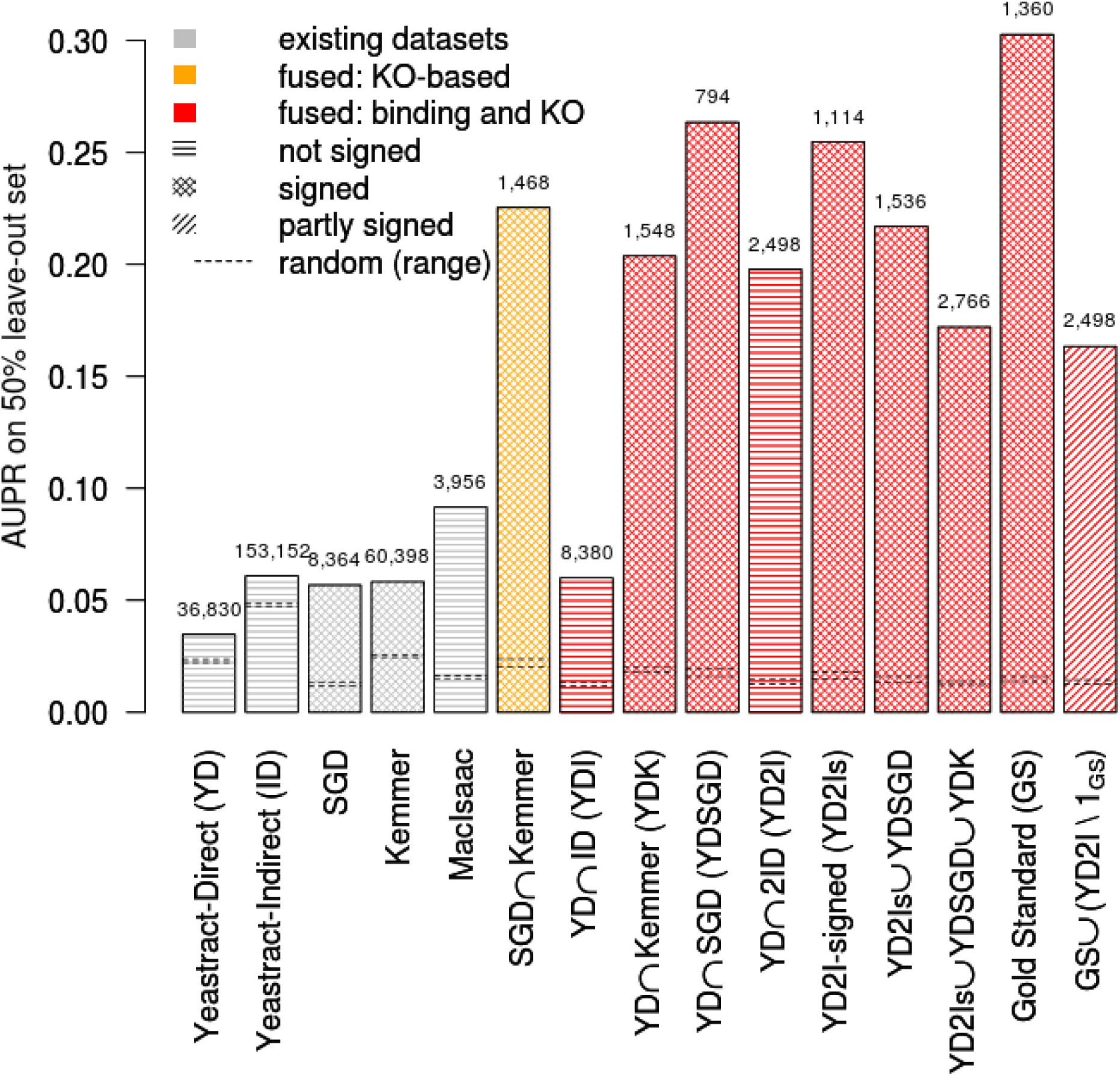
The Gold Standard outperforms other ways of combining the commonly used collections of interactions. Each collection of interactions was split into two equal parts, one for training the Inferelator and one for validation. The height of each bar represents the AUPR calculated on the 50% leave-out set of the corresponding collection of interactions. The number above each bar indicates the number of interactions in the collection, after removing genes and TFs that do not appear in our RNA expression dataset. The color of each bar represents whether it is an existing dataset or a combination of these datasets, either based solely on knock-out evidence (orange), or on an intersection of knock-out and a ChIP binding evidence (red). The shading pattern represents whether none, some, or all interactions are signed. Except for GS, signed interactions came from the signed database in the intersection (Kemmeren or SGD). For the partially signed collection (the rightmost bar), which is a superset of the GS, the same signs as in GS were used when available, and all other interactions were set to 1 (1_GS_ the indicator function of GS, i.e. the GS with all of its interactions set to 1). Standard set theory notation was used, with ∩ denoting union, ⋂ denoting intersection, and \ denoting set difference.

#### 6.1.5 Gene Cluster Half-Life Prediction and Validation

The gene cluster that was most prominently enriched in cytoplasmic translation was also the largest cluster (409 genes), and also contained many genes unrelated to translation. Therefore, for predicting RNA half-lives for cytoplasmic translation genes, we used only the genes that were annotated as “cytoplasmic translation” genes in Saccharomyces Genome Database (GO:0002181). This consisted of 171 unique gene names, 127 of which were ribosomal protein coding genes (either RPLs or RPSs). These genes were used to calculate the predicted translation gene RNA half-lives in Figure 6 (see Section 2.6). The list of these genes can be found in Supplementary File SuppData1.zip.

The predicted RNA half-life values for nucleobase containing small molecule (NCSM) metabolism genes were obtained using the genes in the “nucleic acid metabolism” gene cluster in “log-phase growth” and “chemostat” condition clusters. We used the AmiGO 2 website (Carbon et al., 2009) to obtain 207 genes contained in the nucleobase containing small molecule metabolic process category (GO:0055086). These were the genes for which the experimentally measured RNA half-lives were reported in Figure 6 (Neymotin et al., 2014).

#### 6.1.6 Inferelator Worfklow

The Inferelator is a multi-step framework that aims to predict a TRN from expression data (Bonneau et al., 2006; Greenfield et al., 2013; Arrieta-Ortiz et al., 2015). As inputs, it takes RNA expression data as well as orthogonal sources of data. This orthogonal data can originate from Pol II binding, DNA accessibility, knock-out, overexpression assays, or motif data, and is normally compiled into a matrix of Gold Standard (GS) of interactions or prior known interactions, to be used for training and validation of the resulting network. The main idea behind the Inferelator is that the rate of change of RNA levels of a gene i can be expressed as a difference between its transcription and degradation of its transcript, where the transcription term is approximated as a linear combination of the activities of the gene’s potential TFs. This section describes the details of how we implemented the Inferelator which were not described in Section 2.4.

##### 6.1.6.1 Estimation of Transcription Factor Activities

We calculate TF activities (TFA) of a given TF from the expression levels of its prior known targets in the respective conditions, as described in (Liao et al., 2003; Arrieta-Ortiz et al., 2015). The primary motivation for calculating TFA, rather than using a TF’s RNA expression levels as a proxy for its activity, is that the expression level of a TF is a poor proxy for its activity, and the activity is much better estimated by examining the expression level changes of a TF’s known targets, as demonstrated in Arrieta-Ortiz et al. (2015). We calculate TFA by expressing the response variable, as defined in Equation 2, as a linear combination of TFA levels of each of its known regulators. Mathematically, this relation is expressed as such:

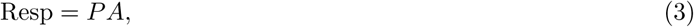

where P is the connectivity matrix, A is the full TFA matrix (with rows corresponding to TFs and columns to samples), and Resp is the response matrix with each row corresponding to the response variable as de ned in Equation 2. The elements of the connectivity matrix P come from the GS or its subset, such that a value of 0 corresponds to no known interaction, 1 to a known positive interaction (activation), and -1 to a known negative interaction (repression). We calculate the activity matrix A by multiplying both sides by the pseudoinverse of P.

##### 6.1.6.2 Model Selection and Prediction Con dence Score

Selecting the set of TFs that regulate genes involves several steps and is the essence of TRN inference, because this is the process in which we select the most likely regulatory network for every gene. The first step involves narrowing down the list of all genes known to be acting as TFs in the organism (in this paper, we use all genes labeled as "transcription factors" or "DNA-binding genes" in SGD as well as all regulators in the YEASTRACT collection of interactions), to a small set of p TFs that are specific to a given gene. This step is completed using time-lagged Context Likelihood of Relatedness (tlCLR), a method for calculating context-dependent mutual information between gene i and its potential TF (Greenfield et al., 2010; Madar et al., 2010). Typically when using the Inferelator, this value is set to p = 10, which is what we use for this paper as well. In addition to these p regulators, TFs known to regulate gene i from prior known interactions are appended to its set of potential regulators. Denote this set as P_i_.

Selecting the most accurate model of regulation for a gene i is thereby equivalent to selecting the set of regulators 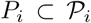 that optimizes an objective function. The choice of the objective function is a version of the Bayesian Information Criterion (BIC), modified with a Zellner’s g-prior, which incorporates prior known interactions into the model by reducing the sparsity penalty for prior known regulatory interactions. This approach, known as Bayesian Best Subset Regression (BBSR), is described in more detail in Greenfield et al. (2013). Note that for this step, we linearly scaled and shifted every gene in the response matrix and every TF in the design (TFA) matrix such that every row has mean 0 and variance 1 across samples, which was necessary in order to avoid having to estimate an extra y-intercept parameter corresponding to basal transcription rate.

Once the model P_i_ that minimizes the BIC is selected, we employ a computational knock-out approach to calculate the confidence we have in each predicted regulatory interaction. The confidence score of the predicted interaction between gene i and TF j ϵ P_i_ is determined in the following manner. We first determine the parameters β_i;k_ for all k ϵ P_i_ by performing linear regression on Equation 2, and use them to calculate the right-hand side of Equation 2. Denote this as the predicted profile. Accordingly, the left-hand side of Equation 2 is denoted as the observed profile. Let σ^2^_i,j_ be the variance of residuals between the predicted profile and the observed profile. Let P_i_^⇁j^ be the same model of regulation of gene i, but without TF j. Then let P^2^_i,⇁j_ be the variance of residuals between the observed profile and the predicted profile as calculated using P^2^_i,⇁j_. Then the confidence score of the interaction between TF j and gene i is de ned as

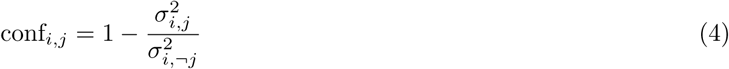

To avoid overfitting and derive empirical confidence intervals for model parameters, tlCLR and BBSR are repeated on different but possibly overlapping subsets of the response matrix, denoted as bootstraps of the expression data. Every such bootstrap is a column-wise (i.e. sample-wise) subset of the response matrix. The columns are selected randomly using the R function sample with default settings. Using this command, we sample n columns with replacement, where n is the total number of columns in the full response matrix. For each bootstrap of the response matrix, the con dence score of every possible interaction is determined using Equation 4. The final combined confidence score for every interaction is determined by rank-combining (adding the ranks of) the confidence scores of that interaction across bootstraps. We use 50 bootstraps for all analyses in this paper, except when otherwise specified.

##### 6.1.6.3 Estimating Network Prediction Accuracy

We define *validation set* as the set of known true interactions that we validate our prediction against. Because the output of the Inferelator is a list of predicted interactions, ranked by their combined confidence scores from highest to lowest, we may define precision and recall as functions of i in the following way: precision is the fraction of the interactions in the top i highest-ranked predicted interactions that are also in the validation set, whereas recall is the number of interactions in the top *i* highest-ranked predicted interactions divided by the total number of interactions in the validation set. We calculate precision and recall for every value of *i* in the full list of predicted interactions. Every PR curve is plotted by connecting the precision and recall values for consecutive values of i with straight lines, and the reported Area Under Precision-Recall curve (AUPR) is the area under this curve.

True Positives (TPs) is a function of *i* that maps *i* to the number of interactions among the top i highest-ranked predicted interactions that are also in the validation set, whereas False Positives (FPs) is the number of interactions among the top i highest-ranked predicted interactions that are not in the validation set. We plot all TP-FP curves by connecting the TP and FP values for consecutive values of *i* with straight lines. We report Area Under the ROC curve (AUROC) by dividing the area under the TP-FP curve by the product of the total number of TPs and the total number of FPs.

For calculating precision and recall, as well as TPs and FPs, on a *leave-out set* (i.e. when a certain *leave-in set* of interactions was used for training the network prediction algorithm, i.e. the TFA step and the BBSR step), we use the leave-out set as the validation set for calculations described above, and the confidence scores of predicted interactions that appear in the leave-in set are set to 0, effectively moving them to the end of the ranked list of predicted interactions. The sign of the interactions is not taken into account in any of our AUPR and AUROC calculations. Predictions involving TFs and genes with no prior known interactions in the leave-out set are excluded from AUPR and AUROC calculations.

#### 6.1.7 Methods Comparisons

For implementing Genie3 (Huynh-Thu et al., 2010), we used the R code downloaded from the Genie3 website (http://www.monteore.ulg.ac.be/huynh-thu/software.html) on September 22, 2016. We modified the original R code to allow for parallel processing, such that calculations on different genes could be performed on separate processors simultaneously. The implementation of the code was conducted as instructed in the README file downloaded from the same website. The R code for iRafNet (Petralia et al., 2015) was downloaded from the iRafNet website (http://research.mssm.edu/tulab/software/irafnet.html) on January 27, 2016. We implemented it using the parameter specifications recommended in the original iRafNet paper (Petralia et al., 2015). In particular, we built the iRafNet weight matrix W using our prior known interactions (the leave-in portion of the Gold Standard) by setting all values of 0 in our leave-in to 0.1 in the weight matrix, and all values of -1 and 1 to 1.1. We set the number of trees ntree to 1000, and number of potential regulators to be sampled from every node to mtry=round 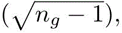 where n_g_ is the number of genes, and round is the rounding function.

For the “Clustering” (no half-life fitting) approach in Table 1 and Table S2, we set τ_i_ = 20 for all genes i and all four condition clusters (see Equation 2 for a definition of τ_i_), which roughly corresponds to an assumed half-life of 14 minutes. This number was based on the median experimentally measured RNA half-life from various recent studies (Neymotin et al., 2014; Munchel et al., 2011; Miller et al., 2011). For the "Fitting" (but no clustering) approach, we used the optimal value of RNA half-life as determined by maximizing the AUPR on the GS-fit collection of leave-out interactions from Split B approach (Figure 1), using the entire RNA expression data set as the input, rather than doing this analysis separately for each condition cluster.

For each of these methods, training was done on the same set of 20 re-samples of the Gold Standard, as described in Sections 2.5 and 2.6, using the Split B aproach, and the AUPRs and AUROCs were calculated using the GS-validate set using the same 20 re-sampes (Figure 1). For each re-sample, individual comparisons in terms of AUPRs and AUROCs between different methods were made, and the number of times, out of 20, that the AUPR (or AUROC) of one method was higher than the other is shown in Table 1 (in terms of AUPR) and Table S2 (in terms of AUROC).

Note that the calculation of RNA half-lives differs between the Split A and Split B approaches. The Split A approach reports a single value for each bi-cluster, which is obtained by taking the median of optimal half-lives across the 20 re-samples, where the optimal half-life is the one that maximizes the AUPR calculated on the GS-fit that corresponds to the given re-sample. Taking the median predicted half-life reduces half-life prediction error due to noise and re-sample specificc effects. In the Split B approach, each RNA half-life prediction is derived from only one GS re-sample, because we prioritized guaranteeing the statistical significance of comparisons between methods in terms of network inference accuracy over RNA half-life prediction accuracy. That is, comparisons between methods were made for each GS re-sample separately, which resulted in 20 points of comparison for each pair of methods, but the method that simultaneously inferred RNA half-life for each bi-cluster did so by maximizing AUPR over GS-fit produced from only one re-sample of the Gold Standard, in contrast with the median over 20 re-samples in the Split A approach. Therefore, the magnitude of the increase in inference accuracy due to half-life fitting that we calculate using the Split B approach (Table 1) is an underestimate of the actual increase in accuracy of our final predicted network, which is produced using the Split A approach.

All of the Inferelator and Genie3 runs that involved the 20x re-sampling of the Gold Standard were performed on the NYU High Performance Cluster (https://wikis.nyu.edu/display/NYUHPC).

## 6.2 Supplementary Figures and Tables

**Figure S4:**
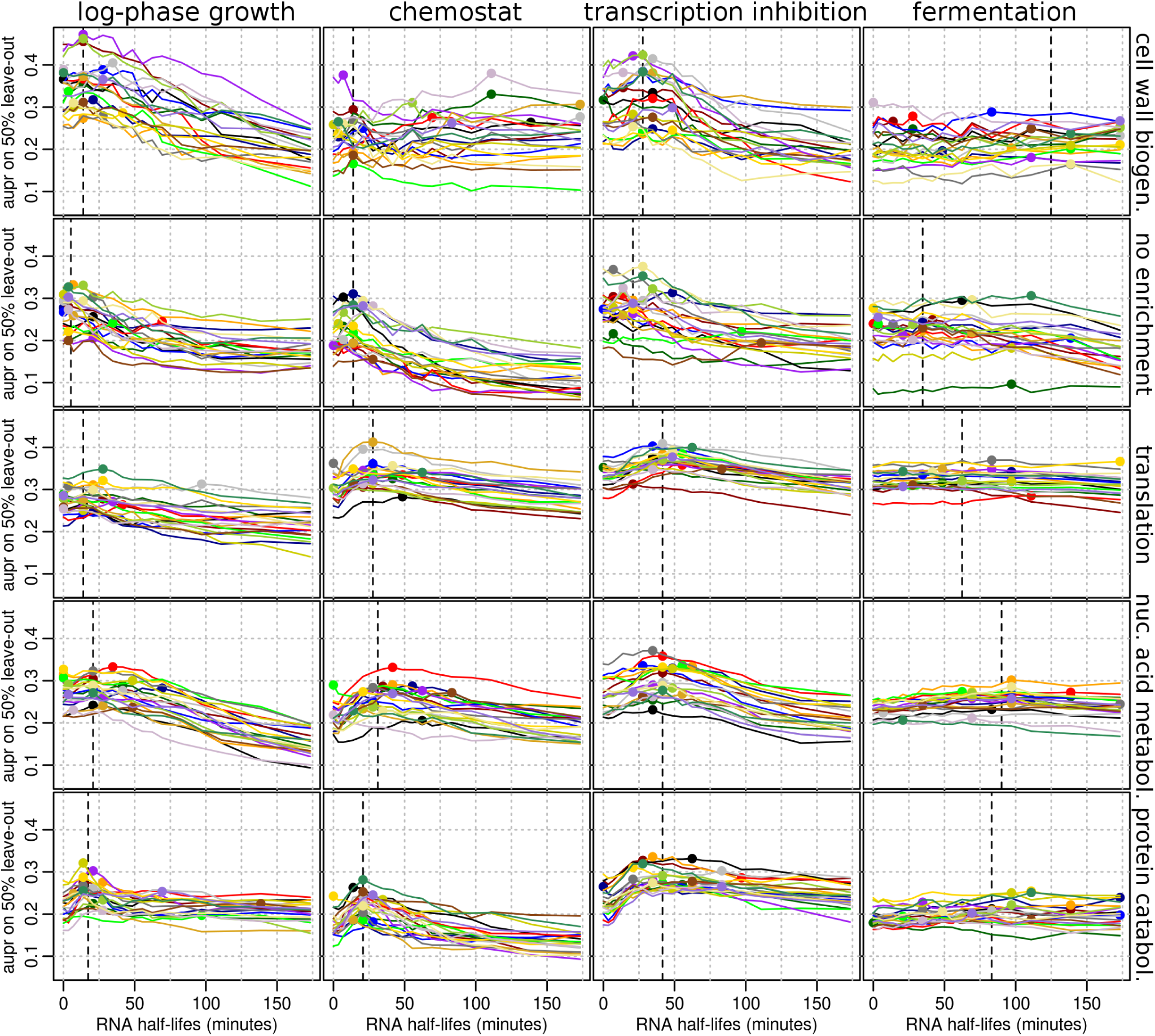
Network inference accuracy is sensitive to RNA half-lives in a condition-specific manner. Each panel shows 20 AUPR plots as as a function of pre-set RNA half-life, when the Inferelator is trained on the the genes specific to the gene cluster (shown on the right) and on conditions specific to the condition cluster (shown on top). Each of the 20 lines corresponds to the 20 Gold Standard re-samples, where 50% of the GS was used for training, and the remaining 50% for calculating AUPR.

**Figure S5:**
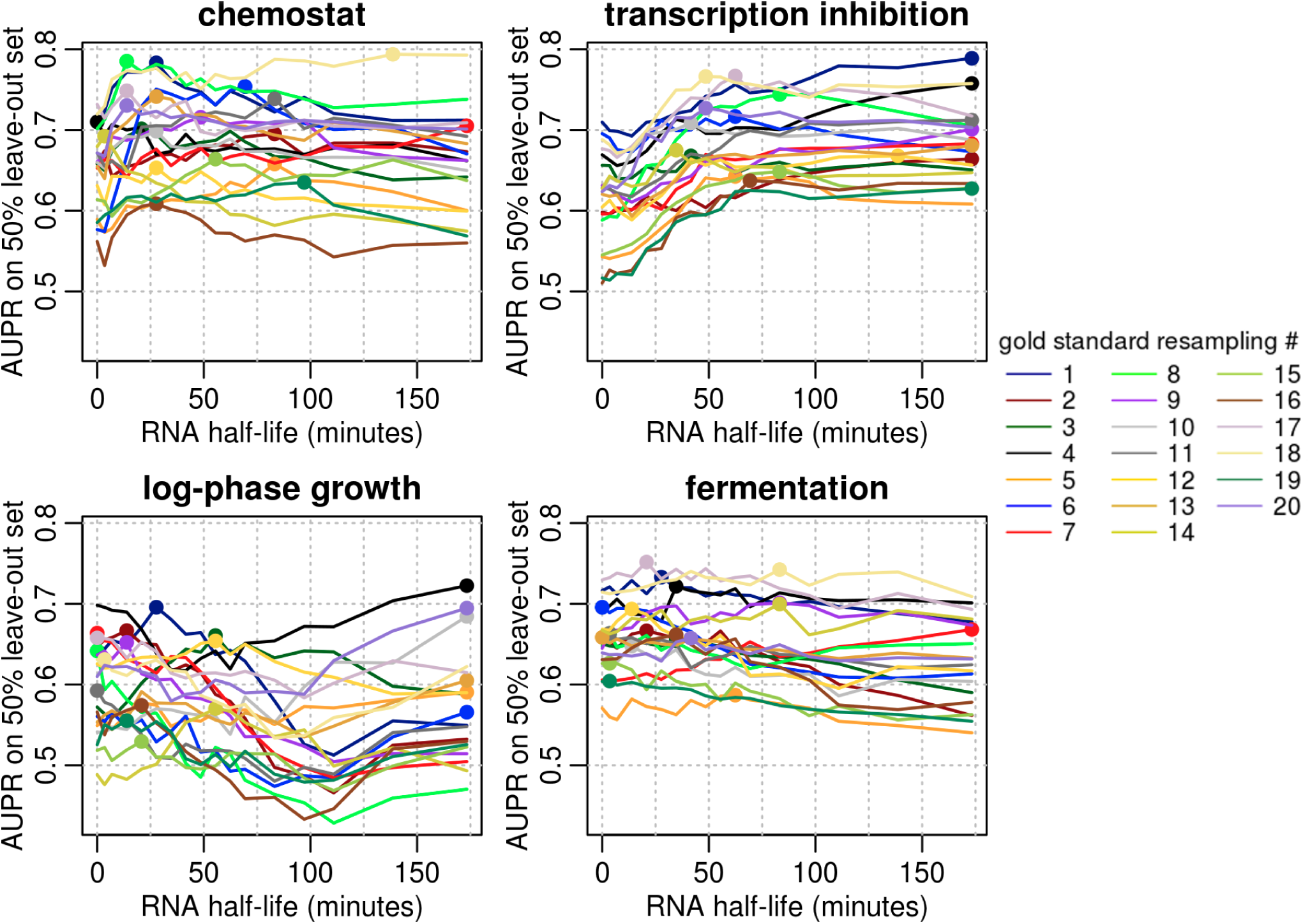
Network inference accuracy is sensitive to RNA half-lives in a condition-specific manner for translation genes. For each cluster, 169 known interactions with 115 "cytoplasmic translation" genes were used to calculate AUPR as a function of RNA half-life. Bold dots correspond to the optimal RNA half-lives for each GS re-sample.

**Figure S6:**
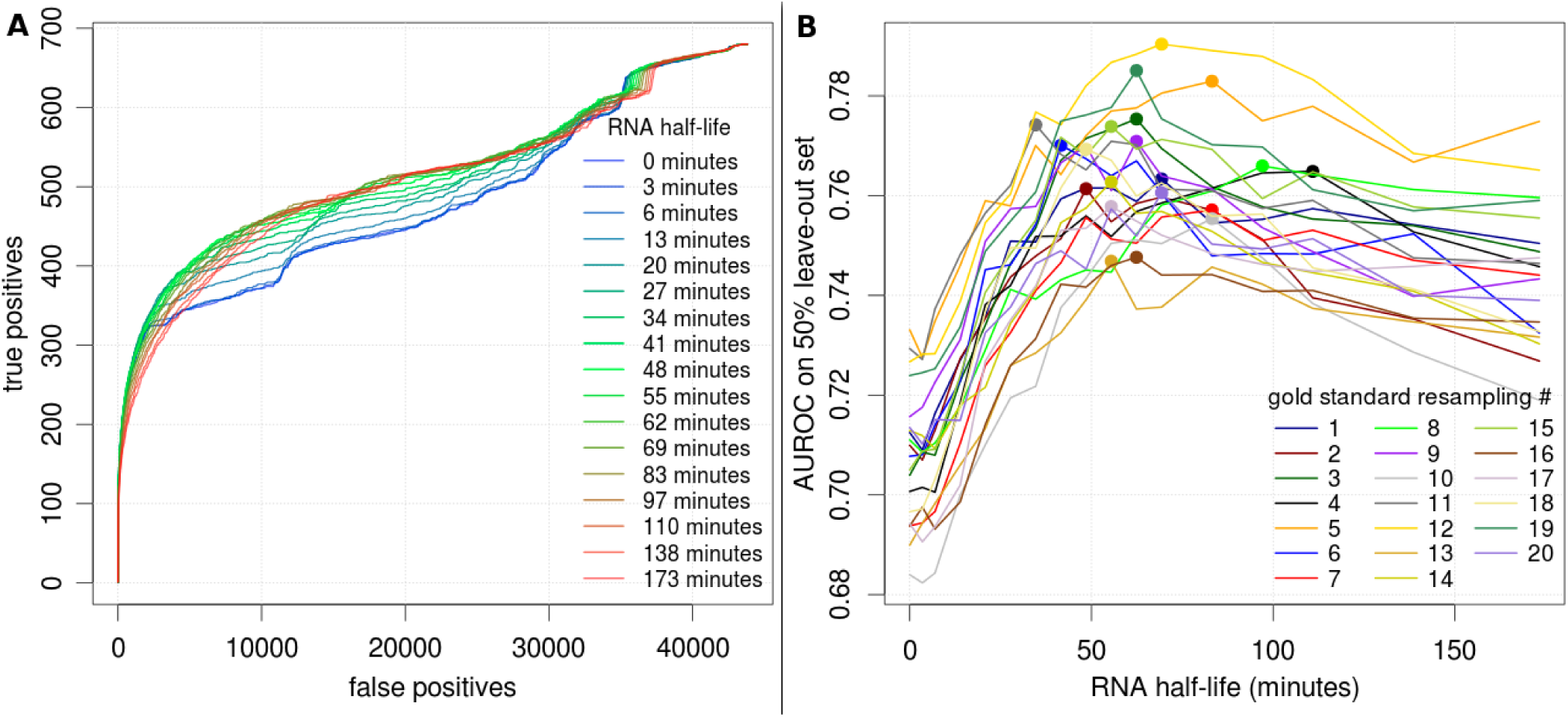
Network Inference is sensitive to RNA half-lives in terms of AUROC. A) True positives vs. false positives curves on the Inferelator output, with each line corresponding to a different pre-set value of RNA half-life. Each line displays the median number of true positives and true negatives across 20 GS re-samplings. B) shows AUROC as a function of pre-set RNA half-life. Different lines denote 20 independent GS re-samplings, and colored dots represent the maximum AUROC for a given GS re-sample.

**Figure S7:**
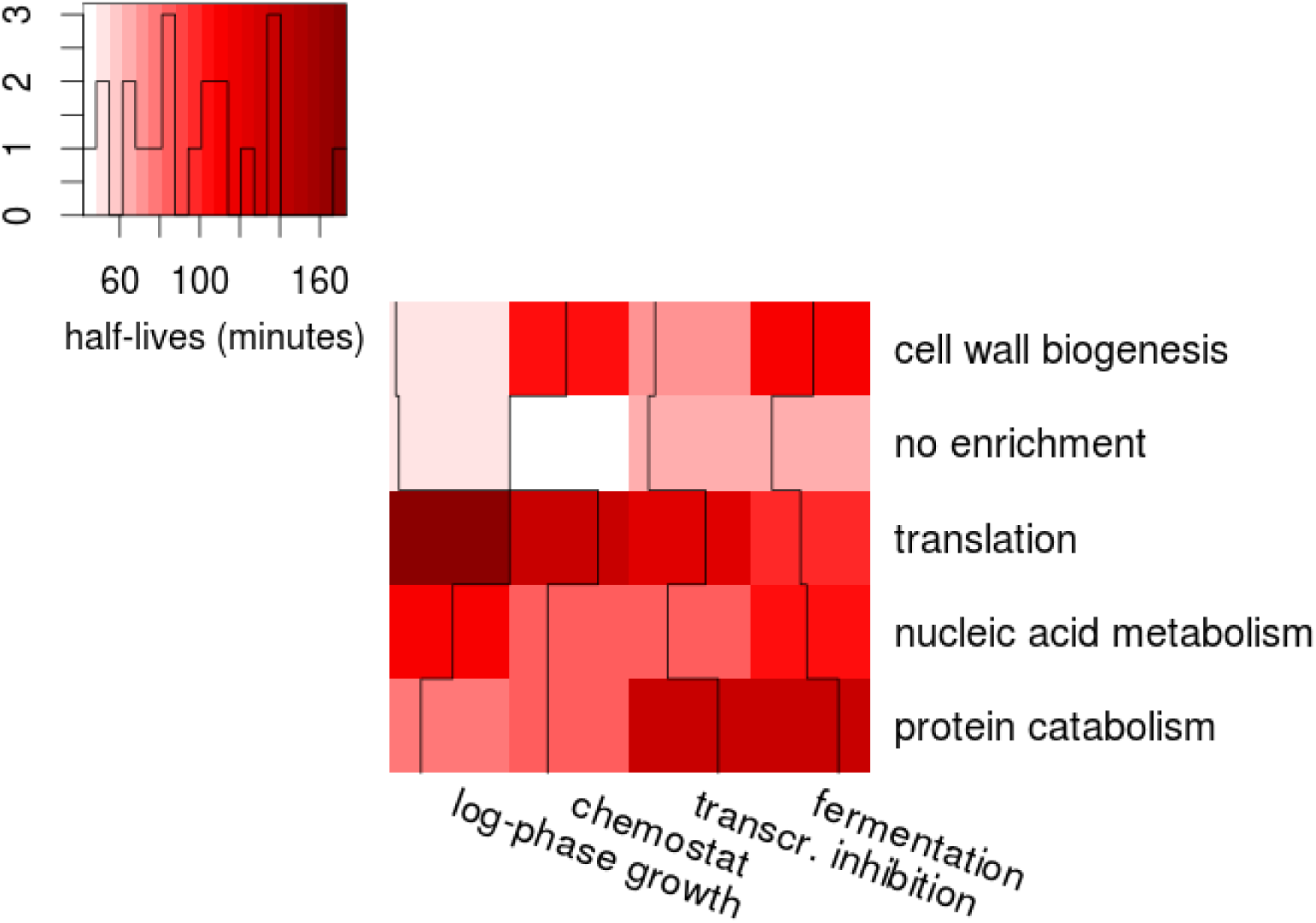
A heatmap of median RNA half-lives that maximized network inference performance with respect to AUROC, for every bi-cluster. This is a summary table of Figure S8.

**Figure S8:**
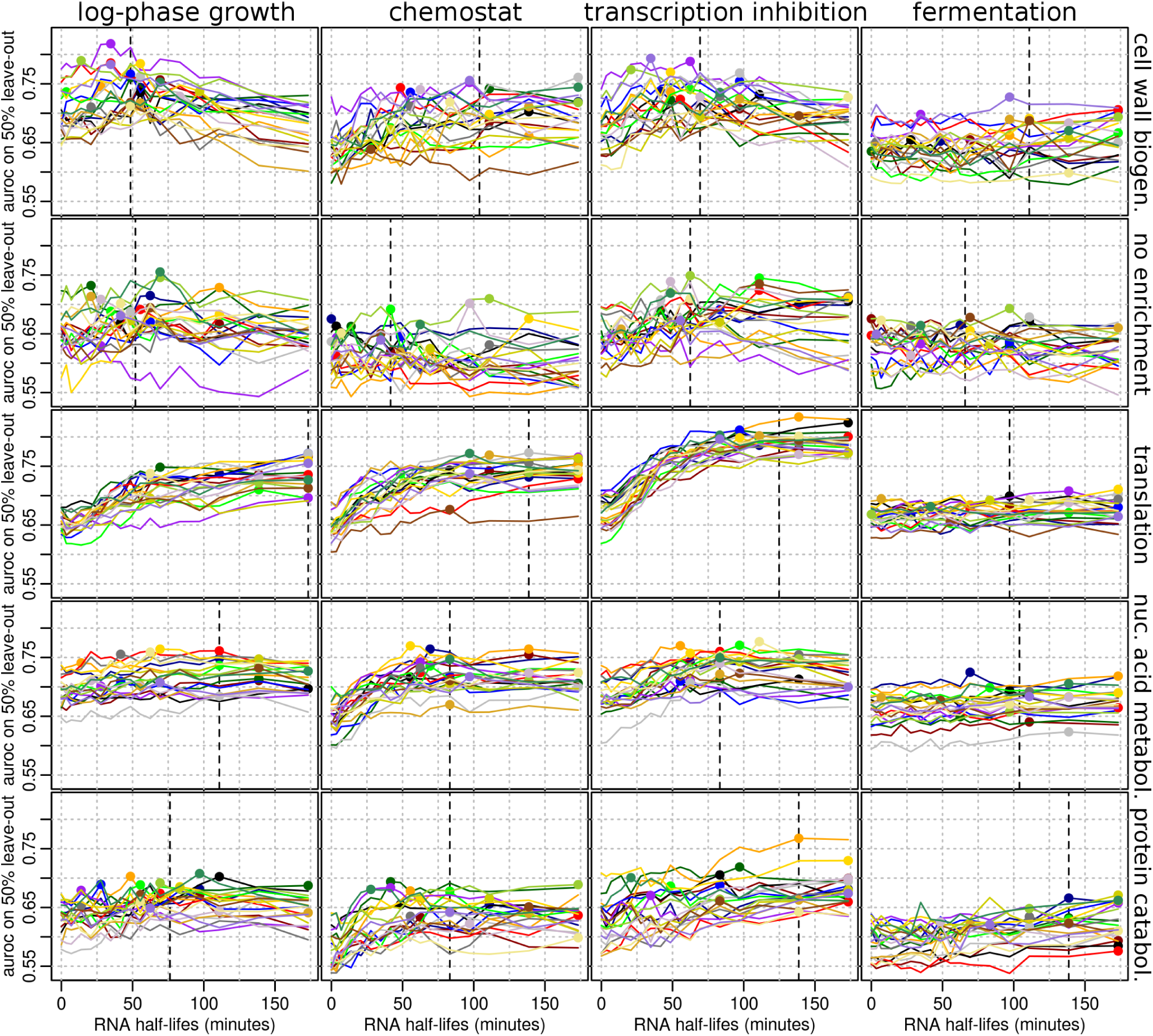
Network inference accuracy is sensitive to RNA half-lives in a condition-specific manner, in terms of area under the ROC curve (AUROC). Each panel shows 20 AUROC plots as as a function of pre-set RNA half-life, when the Inferelator is trained on the the genes specific to the gene cluster (shown on the right) and on conditions specific to the condition cluster (shown on top). Each of the 20 lines corresponds to the 20 Gold Standard re-samples, where 50% of the GS was used for training, and the remaining 50% for calculating AUROC.

**Table S2:**
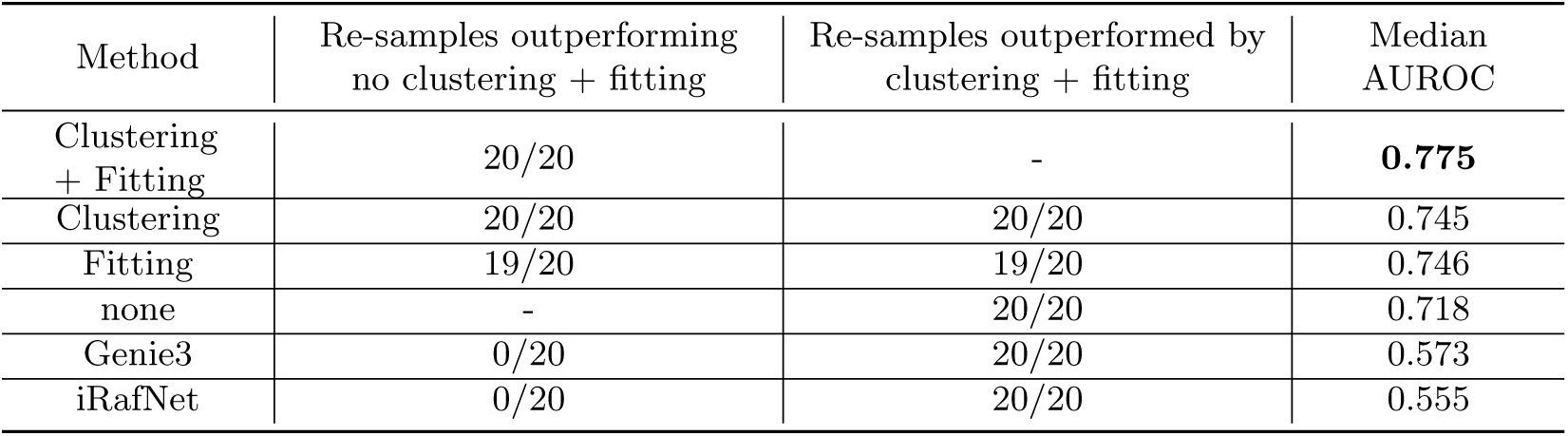
Both expression data bi-clustering and RNA half-life fitting independently improve Inferelator performance (second column). Furthermore, combining the two modifications improves performance as compared to using each of them separately (third column). Columns 2 and 3 show the number of times the AUROC measured on GS-fit from the same re-sample was higher for one method than the other, given a pair of methods specified by the row and the column. Median AUROC for each approach is reported in the fourth column. See Sections 2.5, 2.6, and Sections 6.1.7 for further details.

**Table S3:**
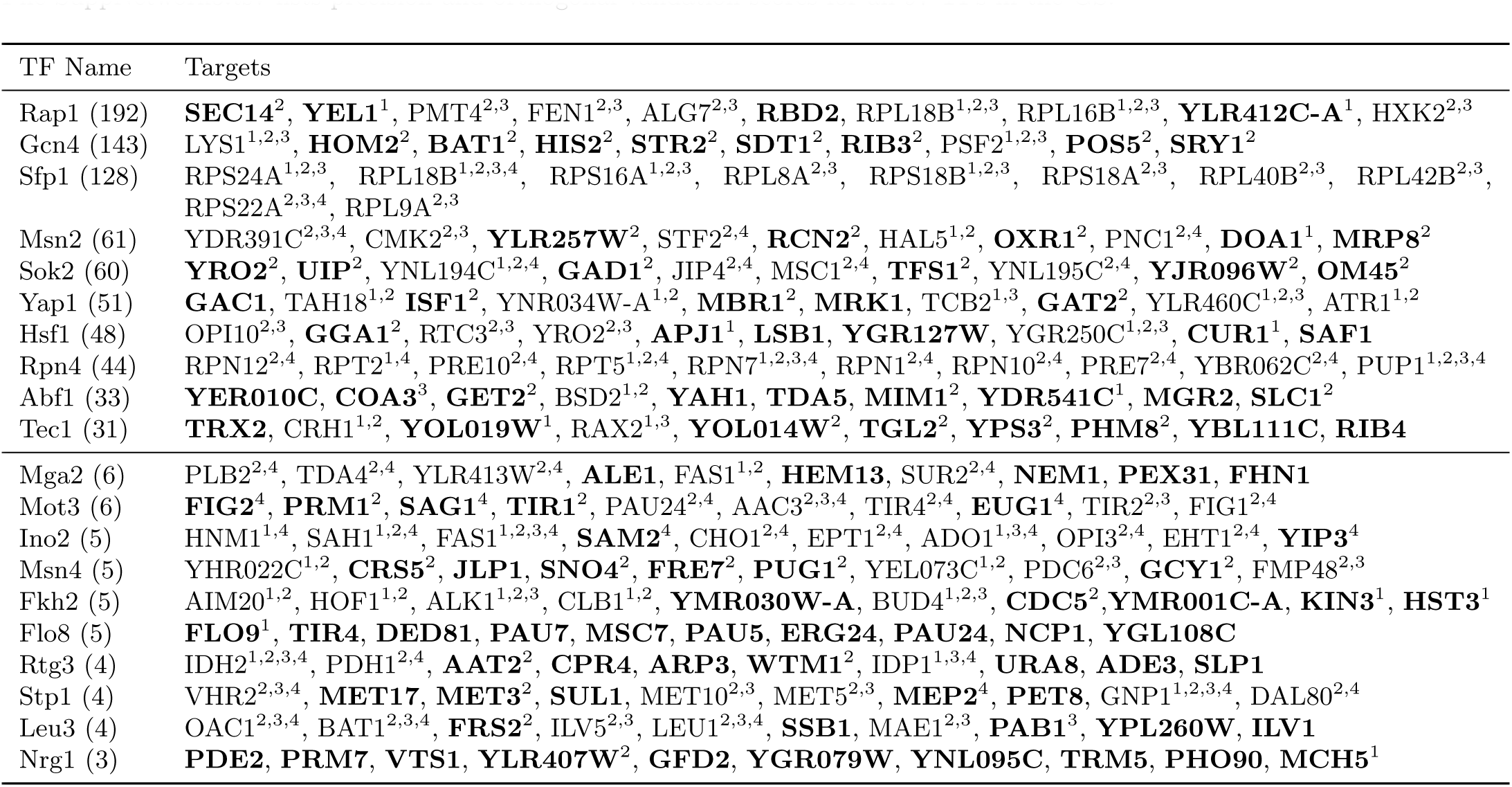
Top 10 new predicted targets for various TFs. The top half of the table lists the 10 TFs that have the highest number of targets in the Gold Standard. For each TF, the number of known targets in the GS is shown in the parentheses, and the top 10 most likely new (i.e. not in the GS) targets are listed. The bottom half displays the median 10 TFs in terms of the number of known targets, out of the 97 TFs with at least one target in the GS. For each TF, targets are listed in the order of decreasing prediction confidence. Superscripts denote whether a given interaction was also present in any of the four large-scale collections of interactions: Yeastract-Direct (1), Yeastract-Indirect (2), SGD (3), and Kemmeren et al. 2014 (4). Bold targets correspond to interactions that are seen in at most one of these four data bases, highlighting completely new interactions. Table 2 displays the precision-based confidence ranks of these interactions, and Supplementary File SuppNetwork3.tsv lists precision and orthogonal validation scores for all 97 TFs in the GS.

**Figure S9:**
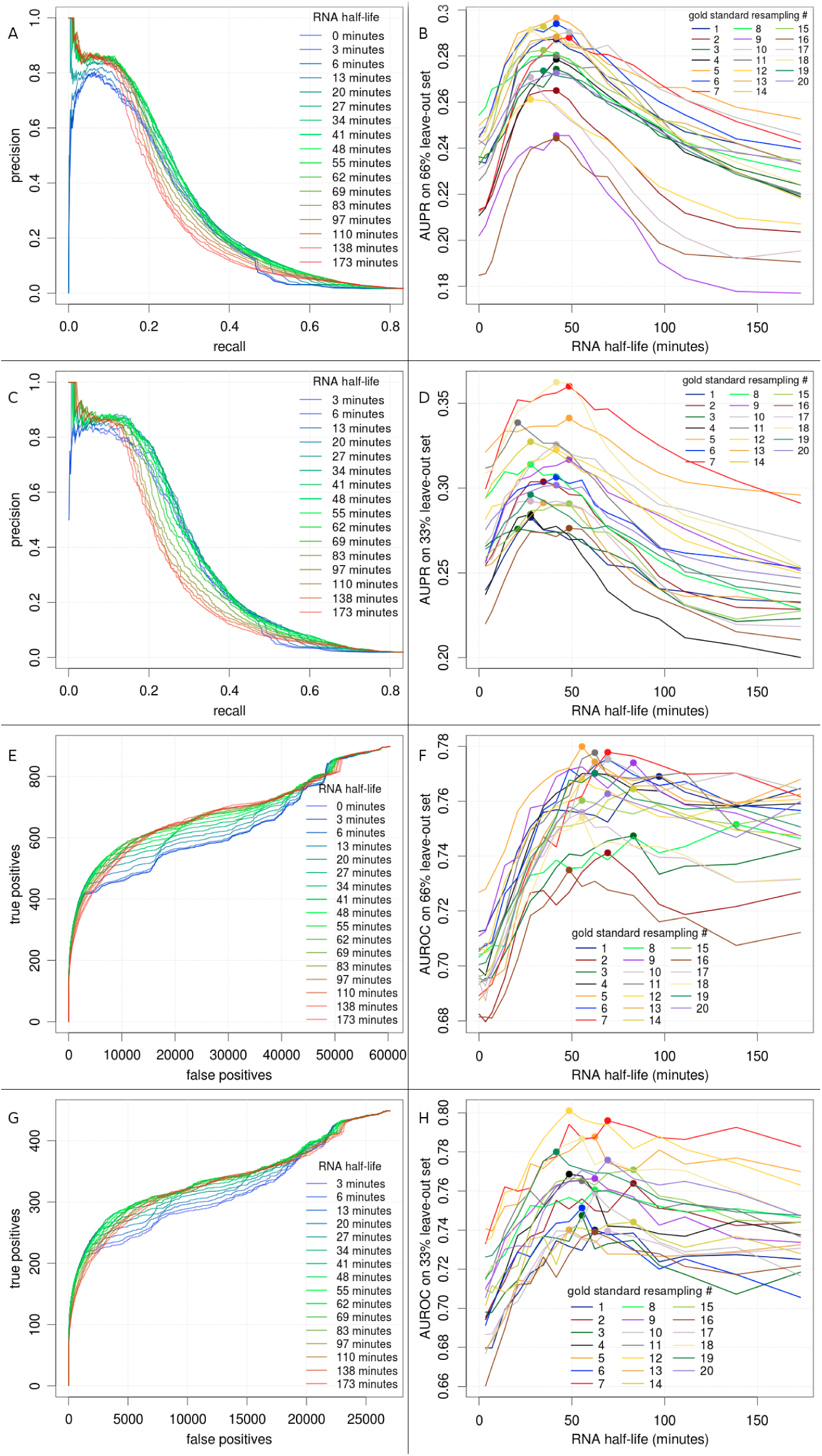
Network inference accuracy is sensitive to RNA half-lives in terms of AUPR and AUROC, independently of the percentage of the Gold Standard used for training. A) and B) The precision-recall curves and the AUPRs as a function of RNA half-life, respectively, for the regime in which 34% of the GS was used for training the Inferelator, and the rest - as a leave-out for validation. C) and D) The same metrics but for a regime in which 67% of the GS was used for training, and the rest for validation. E) and F) The true positives vs. false positives curves and the AUROCs as a function of RNA half-life, respectively, for the regime in which 34% of the GS was used for training, and the rest - for validation. G) and H) The same metrics as E and F, but with 67% of GS as the training set.

## 6.3 Supplementary Data

**SuppNetwork1.tsv** is the final network of predicted interactions at a 0.5 precision cutoff. Each row corresponds to a target gene, each columns corresponds to a TF, and a value of 1 represents a predicted regulatory interaction between the TF and the target gene, whereas a value of 0 represents a lack of regulatory interaction.

**SuppNetwork2.tsv** is a list of all possible TF-gene pairs, ranked in the decreasing order of confidence that there is a regulatory interaction between the pair, according to our final network prediction. The first and the third column list the TF, using its Systematic name and Common name, respectively. The second and fourth columns list the target gene (OLN), using Systematic name and Common name, respectively. The fifth column lists precision value of predictions up to the corresponding interaction, calculated using the Gold Standard (GS). The sixth column represents an interaction’s value in the GS (1 if activating,-1 if repressing, 0 if neither or unknown). Columns 7-10 represent the interaction’s value in Yeastract-Direct, Yeastract-, SGD (all values, not signed), and Kemmeren, respectively.

**SuppNetwork3.tsv** lists the top 1000 newly predicted targets (i.e. these interactions are not in the GS) for the 97 TFs that have known targets in the GS. The first column for each TF lists the targets, the second column lists the precision value of the corresponding TF-target interaction, and the third column represents ":"-delimited numbers with the value of this interaction in Yeastract-Direct (YD), Yeastract-Indirect (YI), SGD, and Kemmeren (K), respectively. For example, "1:0:1:-1" means that the interaction was present in YD, SGD, and Kemmeren, but not YID, and furthermore that the interaction was marked as a repression event in the Kemmeren colleciton of interacitons.

**SuppDoc1.pdf** describes the results of a manual inspection of the Gold Standard targets of the top 15 TFs, listed in the order of decreasing number of known targets in the GS. For each TF, we include the number of positive and negative targets in the GS, known biological attributes of the TF according to SGD, and a discussion of whether the interactions contained in the GS, which were obtained automatically by combining several large data sets, correspond to the established knowledge about the function and the regulation of that TF.

**SuppData1.zip** lists all of the input files required for the inference procedure, including the RNA expression data, the Gold Standard, the meta data, TF names, gene membership and GO enrichments for each gene cluster, cluster membership for each condition cluster, all of the collections of interactions listed in Figure S3, and the list of translation and NCSM metabolism genes used. For more information, see note.txt inside the zip file.

**SuppData2.zip** lists all terms enriched in condition cluster annotation analysis, including a document that describes a manual inspection and con rmation of these automatically-generated assignments. For more information, see note.txt inside the zip file.

**SuppData3.zip** lists all gene-specific and condition-specificτ’s predicted in our analysis. The τ parameter is defined in Equation 2, and can be converted to RNA half-lives by being multiplied by ln (2). The R code is available upon request.

